# Cataloguing and profiling of the gut virome in Chinese populations uncover extensive viral signatures across common diseases

**DOI:** 10.1101/2022.12.27.522048

**Authors:** Shenghui Li, Qiulong Yan, Yue Zhang, Ruochun Guo, Pan Zhang, Qingbo Lv, Fang Chen, Zhiming Li, Jinxin Meng, Jing Li, Guangyang Wang, Changming Chen, Hayan Ullah, Lin Cheng, Shao Fan, Rui Li, Wei You, Yan Zhang, Jie Ma, Wen Sun, Xiaochi Ma

## Abstract

The gut viral community has been linked to human physiology and health, but our knowledge of its genetic and functional contents and disease dependence is far from complete. Here, we collected 11,327 bulk or viral metagenomes from fecal samples from large-scale Chinese populations to establish a Chinese gut virus catalogue (cnGVC) comprising 67,096 nonredundant viral genomes. This catalogue included ∼70% of novel viruses that are not represented in existing gut viral databases, and allowed us to characterize the functional diversity and specificity of the gut virome. Using cnGVC, we 1) profiled the gut virome in large-scale populations and evaluated their sex- and age-related variations, 2) investigated the diversity and compositional patterns of the gut virome across common diseases by analyzing 6,314 bulk metagenomes spanning 28 disease or unhealthy statuses, and 3) identified a large number of universal viral signatures of diseases and validated their predictive ability for health status. Overall, our resources and results would contribute to the grand effort of expanding the knowledge of the human gut virome and addressing a full picture of the associations between viruses and common diseases.

## Introduction

The viral community in our gut, referred to as the human gut virome, is a key component of the gut microbial community with extensive unexplored genetic and functional diversity^1^. Traditional gut virology research usually focused on some specific enteroviruses^2, 3^, but its results and applications were very limited due to the insufficiency of virus discovery. However, with the rapid development of high-throughput whole-metagenome (bulk) sequencing and virus-like particle (VLP)-based virome sequencing technologies, some characteristics of the gut virome have been preliminarily delineated. Generally, the normal gut viral composition is highly individual-heterogeneous and depends on human sex, age, geography, and lifestyle^4–7^. Longitudinal analysis revealed that the gut virome of healthy adults is temporally stable but highly variable along with the changes in the external environment^4^. Significant alterations of the gut virome have been observed in a diverse range of gastrointestinal and systemic disorders, including colorectal cancer (CRC)^8, 9^, inflammatory bowel disease (IBD)^10^, necrotizing enterocolitis^11^, liver disease^12, 13^, autoimmune disease^14^, metabolic syndrome^15^, or even infectious diseases such as acquired immunodeficiency syndrome (AIDS)^16^ and COVID-19^17, 18^. These efforts suggest a critical role of the gut virome in human health and raise the requirement for evaluation of its variation pattern and pathophysiological role in depth and breadth.

Catalogues of reference viral genomes in the human gut virome are crucial for quantificational and functional analyses^19^. Recent studies identified a tremendous number of viruses from publicly available fecal metagenomes and created the Gut Virome Database (GVD)^6^, Gut Phage Database (GPD)^1^, and Metagenomic Gut Virus (MGV) catalogue^20^ that contained tens of thousands of viruses for each. A surprising phenomenon was that the viral sequences uncovered by different catalogues substantially differed (sharing <50% of each other; Supplementary Fig. 1a). Even in the same catalogue, it showed only a small proportion of viruses are shared between the samples from different regions (Supplementary Fig. 1b). Although technical differences may lead to some, these findings strongly suggested that the gut virome is highly heterogeneous among different populations, as also supported by other studies^7, 21^. With the rapid expansion of bulk and VLP-based virome metagenomic samples, the representativeness of the gut viral catalogue can be significantly improved by using large-scale samples from a single population and the unified state-of-the-art processing pipelines^19^.

To expand the viral genome reference and provide a more comprehensive view of the gut virome, herein, we built a catalogue of the gut viruses by processing over 11,000 fecal bulk or VLP-based viral metagenomes of Chinese origin. The catalogue was named Chinese gut virus catalogue (cnGVC) including 67,096 nonredundant viruses (dereplicated from 426,496 viruses with >95% nucleotide similarity) with a majority never found in existing gut viral databases. The cnGVC improved functional characterization of the gut virome at high resolution, largely increased the recruitment of gut viruses in metagenomes (capturing 19.8% viral reads in bulk metagenomes and 56.7% in viral metagenomes), and enabled to evaluate of the sex- and age-related variations in virome. In terms of disease, we profiled the gut viromes of 6,314 fecal metagenome samples spanning 28 disease or unhealthy statuses and found that a majority of the investigated diseases occurred with significant reductions in viral richness and diversity and remarkable shifts in the overall gut viral composition. Moreover, we identified 7,161 differential viruses using meta-analysis across all diseased and healthy individuals and demonstrated the potential of these universal viral signatures in predicting human health status.

## Results

### Construction of the gut virus catalogue in Chinese populations

To expand the resource for gut virome research, we downloaded and reanalyzed raw data from a collection of 10,170 fecal bulk metagenomes and 1,157 fecal viral metagenomes deriving from 50 previously published studies (Supplementary Fig. 2a; Supplementary Table 1). This dataset contained samples spanning 18 provincial-level administrative regions of China (Fig. 1a), which represents the current largest fecal metagenomic dataset of Chinese populations. These samples contained a total of 92.2 Tbp of high-quality non-human metagenomic data after being processed with a unified pipeline, and further generated a total of 290 million long contigs (≥5 kbp; total length 2.3 Tbp) via de novo metagenomic assembly for each sample. We identified a total of 426,496 highly credible viral sequences (estimated completeness ≥50%) from the contigs using an integrated homology- and feature-based pipeline (see Methods), henceforth referred to as the cnGVC. The length of viral sequences ranged from 5,004 bp to 504,568 bp, with an average length of 37,961 bp and an N50 length of 44,025 bp (Supplementary Fig. 3a). We evaluated the completeness and contamination of these viruses using the CheckV algorithm^22^, revealing 15.7% complete, 31.8% high-, 52.5% medium-completeness viruses (Fig. 1b). And 99.3% of these viral genomes were not contaminated (Supplementary Fig. 3b), meaning that no microbial-specific genes were detected at the terminals of these viruses.

**Fig. 1.**
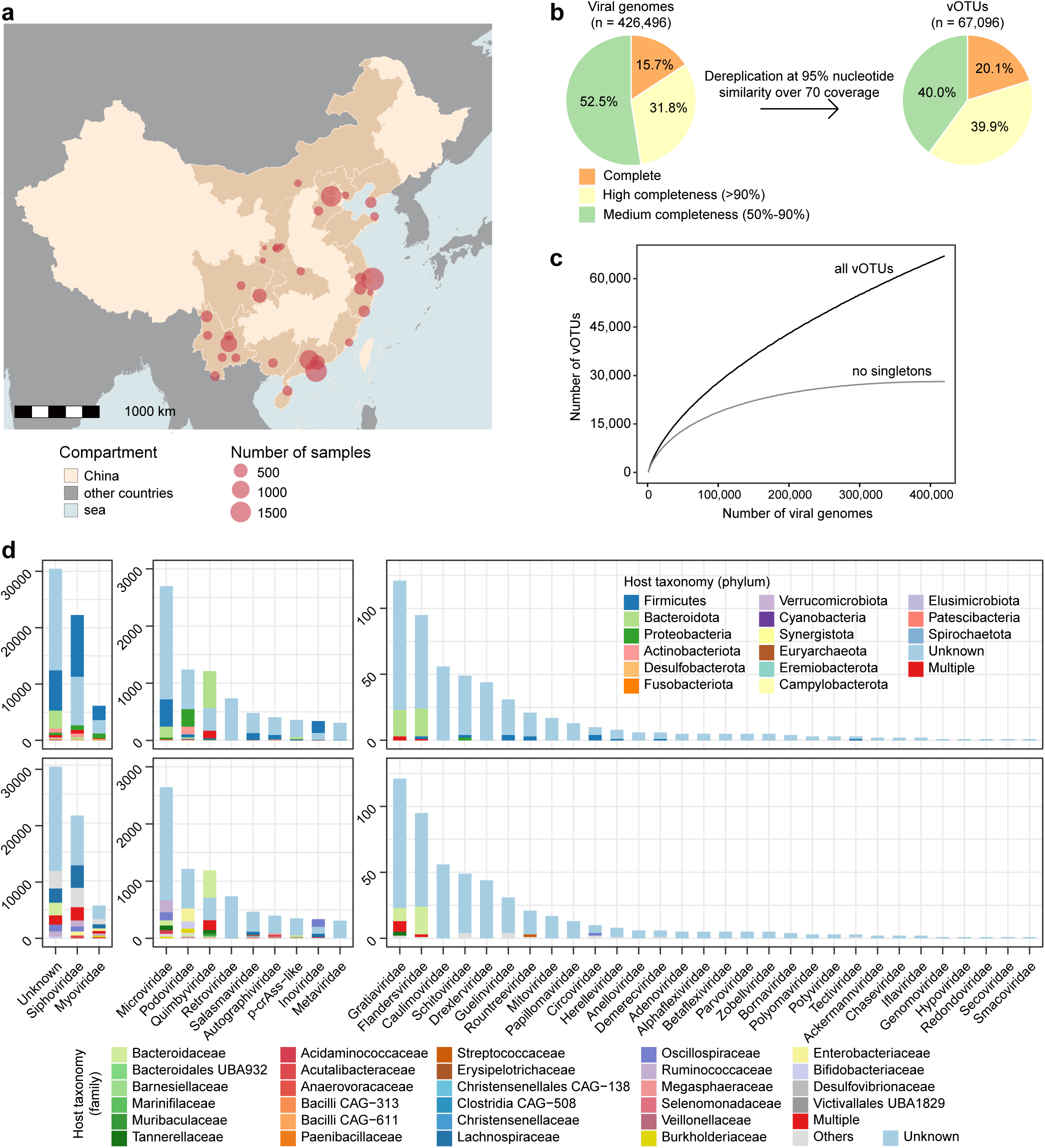
Overview of the cnGVC. **a,** Map of China showing the geographic distribution of metagenomic samples used to construct the gut virus catalogue. **b,** Pie plots showing the estimated completeness of all viruses (left panel) and nonredundant vOTUs (right panel) in cnGVC. **c,** Rarefaction curves of the number of vOTUs and no-singletons as a function of the number of all viral genomes. **d,** Distribution of taxonomic annotation and host assignment of the cnGVC. The vOTUs are grouped at the family level, and the prokaryotic host taxa are shown at the phylum (upper panel) and family (bottom panel) levels. The number of vOTUs that had more than one predicted host is labeled by red color.

Next, we clustered the viral sequences at a 95% average nucleotide similarity threshold (≥70% coverage) and resulted in a nonredundant virus catalogue with 67,096 viral operational taxonomic units (vOTUs). For each vOTUs, the longest viral sequence was selected as the representative virus. 20.1% of vOTUs contained a complete representative virus as estimated by CheckV^22^, while the representative viruses of 39.9% and 40.0% of vOTUs were high- and medium-completeness, respectively (Fig. 1b). Noticeably, only 41.9% (28,132/67,096) vOTUs contained 2 or more viral members, whereas the remaining 58.1% vOTUs were singletons. We estimated the accumulation number of vOTUs as a function of the number of viral genomes to evaluate the coverage of the viral space. The accumulation curve of vOTUs was not yet reaching a plateau, whereas the curve of no-singleton vOTUs appeared to be approaching an asymptote (Fig. 1c). These findings suggest that the viral space of the core gut virome is close to saturation in the current sample size. However, a large number of rare members of the gut virome remain to be discovered.

54.6% (36,651/67,096) of the nonredundant vOTUs could be robustly assigned to a known viral family. In agreement with the previous studies^5, 23^, three families of the double-stranded DNA (dsDNA) Caudovirales order (i.e., Siphoviridae, Myoviridae, and Podoviridae) and the single-stranded DNA (ssDNA) Microviridae family constituted the vast majority of taxonomically assigned vOTUs (Fig. 1d). While the other representatives included Salasmaviridae, Autographiviridae, Inoviridae, and some eukaryotic viruses (e.g., Retroviridae). 357 vOTUs were robustly assigned into the crAss-like viruses and were divided from other Podoviridae members due to their unique genomic features^24^. Three new candidate families, including Quimbyviridae, Gratiaviridae, and Flandersviridae, that were recently identified from the human gut virome^25^, also frequently appeared in our catalogue.

On the basis of similarity to genome sequences or CRISPR spacers in the comprehensive unified human gastrointestinal genome (UHGG) database^26^, 48.3% of the 67,096 vOTUs could be assigned into one or more prokaryotic hosts. The most common identifiable hosts of Siphoviridae, Myoviridae, and Microviridae members were Firmicutes species, but their hosts at the family level were largely differed (Fig. 1d). The major hosts of crAss-like, Quimbyviridae, Gratiaviridae, and Flandersviridae were Bacteroidetes species, and particularly, a large proportion of viruses of the later three families were predicted to infect Bacteroidaceae. Podoviridae viruses were generally predicted to infect Proteobacteria (mainly Enterobacteriaceae) and some Firmicutes species. 6.3% (2,031/32,406) of the annotated vOTUs had hosts that belonged to two or more bacterial phyla, and 13.6% (4,423/32,406) vOTUs had hosts across different families, suggesting a relatively narrow host range of most gut viruses.

### Comparison with existing gut viral databases

We compared the nonredundant viruses of cnGVC with three available human gut viral catalogues (i.e., GVD, GPD, and MGV) and the gut viruses from CHVD (CHVD-gut). All four existing catalogues were filtered to retain high- and medium-quality viruses (estimated completeness ≥50%) and then dereplicated using the same thresholds with cnGVC (95% ANI and 70% coverage). After this processing, the MGV and GPD had the largest number of vOTUs in the existing catalogues, which contained 43,054 and 39,172 vOTUs, respectively; while the CHVD-gut and GVD had only 18,646 and 8,882 vOTUs, respectively (Supplementary Fig. 4). Pooling of all databases revealed that 70.2% (47,118/67,096) of vOTUs in the cnGVC are not found in other catalogues (Fig. 2a). These results indicate that, even in the presence of current multiple gut viral catalogues, there are still a large number of novel viruses in cnGVC, which probably relates to the existence of a large number of Chinese samples in cnGVC. Moreover, we found that the cnGVC covered 47.8% of the vOTUs from the Asia samples of MGV, but only 25.8% of the vOTUs from the non-Asia samples of MGV were covered (Fig. 2b), suggesting the higher representative of cnGVC in Asian rather than Western populations.

**Fig. 2.**
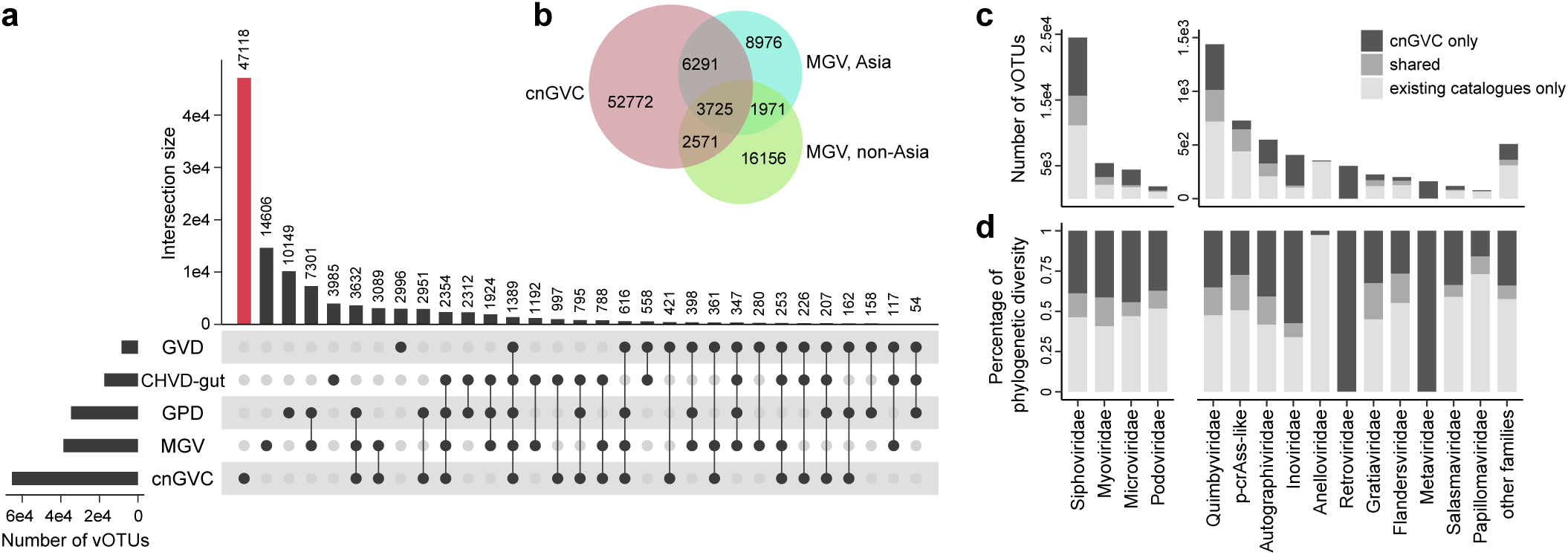
Comparison between cnGVC and other existing databases. **a,** UpSet plot showing the number of vOTUs shared by existing gut viral catalogues. The vOTUs that are uniquely found in cnGVC are labeled by red color. CHVD-gut, gut viruses from the Cenote Human Virome Database; GVD, Gut Virome Database; GPD, Gut Phage Database; MGV, Metagenomic Gut Virus. **b,** Venn plot showing the sharing relationship of vOTUs in the cnGVC and MGV catalogues. Viruses in MGV are divided by their origin in Asia or non-Asia samples. **c,** Comparison of vOTUs between cnGVC and other existing databases at the family level. **d,** Contribution of phylogenetic diversity (PD) by the viruses from cnGVC and other existing databases. Phylogenetic trees are constructed for each viral family, and the PDs are calculated for cnGVC-specific vOTUs, existing-catalogue-specific vOTUs, and shared vOTUs accordingly.

To further illustrate the novelty of cnGVC in the phylogenomic scope, we next compared all high completeness (≥90%) viruses between cnGVC and the existing gut viral catalogues at the family level. For almost all families, cnGVC greatly expanded the content of known high-completeness viruses from the human gut (Fig. 2c). It increased the number of gut-derived vOTUs by 41.8%-112.1% (average 64.8%) for the top five dominant families (i.e., Siphoviridae, Microviridae, Myoviridae, Podoviridae, and Quimbyviridae). 304 high-completeness viruses belonging to Retroviridae were uniquely found in the cnGVC catalogue. Additionally, cnGVC included 160 viruses belonging to Metaviridae, but only 1 high-quality Metaviridae virus existed in the other catalogues. Conversely, vOTUs of the small circular ssDNA viral family Anelloviridae were mainly found in the existing catalogues but rarely occurred in cnGVC. Moreover, genome-based phylogenetic analyses for five dominant families revealed that the newly-found vOTUs in cnGVC are broadly distributed in the major taxonomic lineages across the phylogenetic trees (Supplementary Fig. 5), suggesting that they may fill the gaps in the viral tree of life in the human gut. In addition, we calculated the phylogenetic diversity (PD) based on phylogenetic trees for each viral family and found that the cnGVC-specific viruses accounted for, on average, 39.4% of the PD in the trees for all families (Fig. 2d). Together with the aforementioned findings, our results highlight the high comprehensiveness and novelty of the viral catalogue from large-scale Chinese populations.

### Functional configuration of the gut viruses

Our extended viral catalogue may enable high-resolution functional analysis of the gut virome. For this purpose, we predicted approximately 22.0 million protein-coding genes from the 426,496 viral genomes of cnGVC and clustered them into 1,595,487 nonredundant genes at 90% average amino acid identity (AAI) (Fig. 3a). The nonredundant gene catalogue contained 96.6% of complete genes, which is referred to as the current, and to our knowledge, largest viral gene database of the human gut. Similar to the vOTUs, rarefaction analysis showed that the accumulation curve of no-singleton genes of the viral gene catalogue has approached an asymptote (Fig. 3b).

**Fig. 3.**
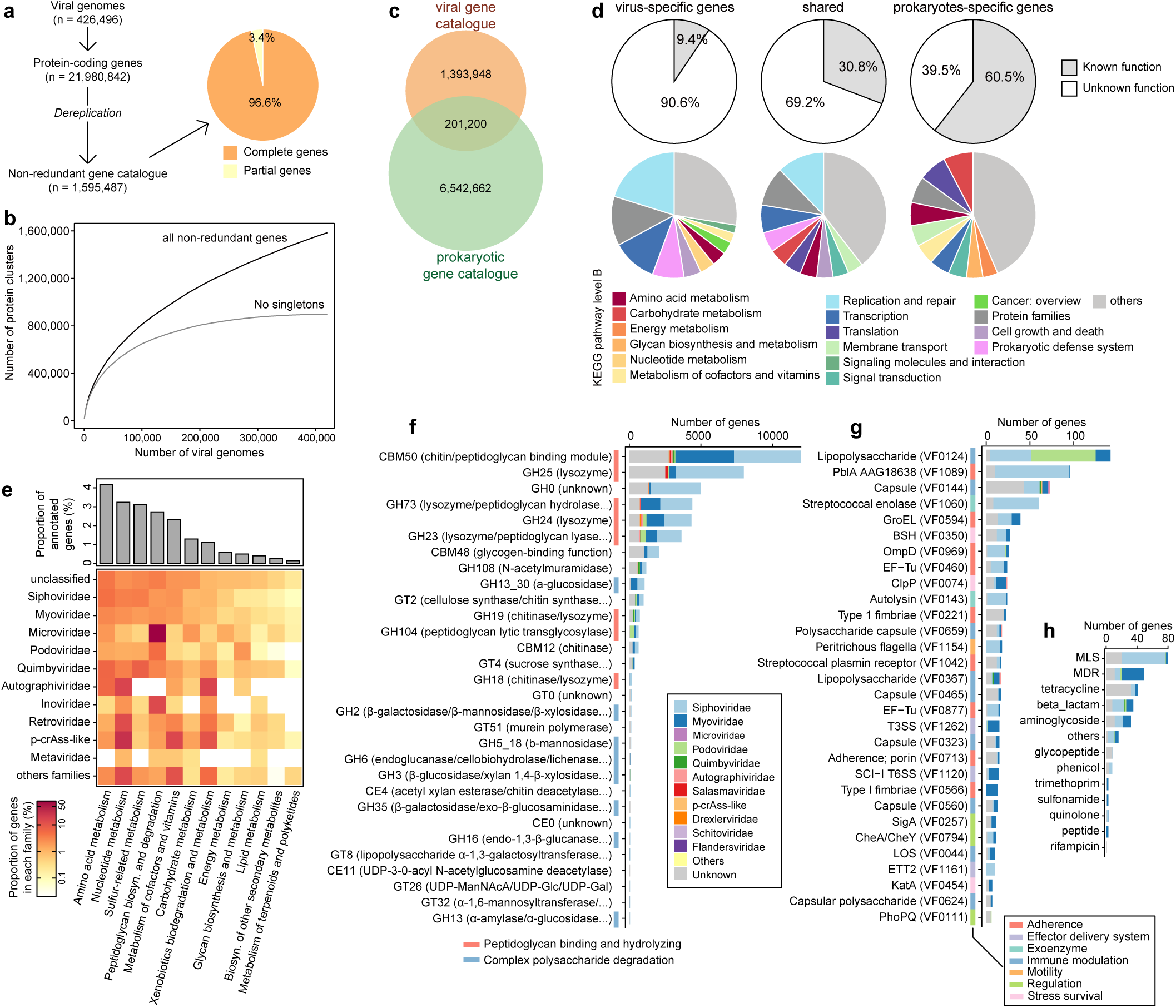
Overview of the viral functions of cnGVC. **a,** Construction of a nonredundant gene catalogue from the cnGVC viral genomes. **b,** Rarefaction curves of the number of nonredundant viral genes and no-singletons as a function of the number of all viral genomes. **c,** Venn plot showing the sharing relationship between gut viral genes and prokaryotic genes. **d,** Functional composition of the virus-specific genes, prokaryotes-specific genes, and shared genes. Functions are categorized at the KEGG pathway level B. **e,** Composition of viral auxiliary metabolic genes (AMGs) for each viral family. AMGs are grouped at the KEGG pathway level B and sulfur-related metabolism. The bar plot (upper panel) shows the overall proportions of AMGs versus the number of annotated genes for all viruses, and the heatmap (bottom panel) shows the proportions of AMGs versus the number of annotated genes for each family. **f-h,** Distribution of virus-encoded carbohydrate-active enzymes (CAZymes) **(f)**, virulence factor genes (VFGs) **(g)**, and antibiotic resistance genes **(h)** for each family. For CAZymes and VFGs, only the top 30 enzymes are shown.

To further demonstrate the functional specificity of the gut virome, we first compared the viral genes with the gut prokaryotic gene catalogue UHGP-90 (the unified human gastrointestinal protein database clustered at 90% AAI)^26^. Although UHGP-90 was generated by an extensive collection of global gut prokaryotic genomes^26^, it covered only 12.6% of viral genes in cnGVC (Fig. 3c). This finding reveals a substantial difference in gene contents between the gut viruses and prokaryotes and also suggests that the gene/functional specificity of the gut virome has been underestimated in the past. However, only 9.4% of the virus-specific genes were functionally known under the Kyoto Encyclopedia of Genes and Genomes (KEGG) database^27^, this proportion was significantly lower than that of the prokaryote-specific genes (60.5%) and shared genes (30.8%) (Fig. 3d). Consistent with the previous studies^20, 22^, virus-specific genes are dominated by the functions involving to replication and repair, transcription, prokaryotic defense system, and metabolism of amino acids and nucleotides. We next focused especially on the viral auxiliary metabolic genes (AMGs), since the virus-encoded AMGs have been recognized to redirect host functional capacities thereby directly influencing the gut ecosystem^28, 29^. Nearly one-fifth (19.8%) of the KEGG-annotated genes, corresponding to 2.4% of all genes, were identified as AMGs based on a previously curated list^30^. Metabolism of amino acids, nucleotides, and sulfur compounds were the most popular auxiliary metabolic functions, and these functions were widely encoded by almost all viral families except for some “small” viruses such as Autographiviridae (absent sulfur metabolism genes) and Metaviridae (absent amino acid and sulfur metabolism genes) (Fig. 3e). Microviridae members had the highest proportion of genes involving peptidoglycan degradation (mainly consisted of the zinc D-Ala-D-Ala carboxypeptidase K08640), while crAss-like viruses had a comparatively high proportion of genes involving to the metabolism of cofactors, vitamins, and xenobiotics.

Finally, we uncovered 48,358 auxiliary carbohydrate-active enzymes (CAZymes) (corresponding to 3.0% of all genes), 983 virulence factor genes (VFGs) (0.062%), and 283 antibiotic resistance genes (ARGs) (0.018%) from the viral nonredundant gene catalogue (Fig. 3f-h). A majority of (>70%) the CAZymes were involved in binding and hydrolyzing bacterial peptidoglycans, which is probably associated with the degradation of peptidoglycans of the bacterial cell wall during infection^31^. Besides, many virus-encoded CAZymes belonged to the glycoside hydrolase families that are involved in the decomposition of complex polysaccharides (Fig. 3f), and a large number of these genes (n = 546; Supplementary Table 2) were also validated with functions in bacterial polymer (e.g., pectin, cellulose, and xylan) hydrolysis by three-dimensional protein structural modeling. These results largely agree with previous studies in environmental viral communities showing the important role of polysaccharide decomposition in viral ecology^32, 33^. Notably, the polysaccharide-degrading enzymes were frequently encoded by members of the Siphoviridae and Myoviridae but not in Podoviridae and Quimbyviridae viruses, despite the latter two families having bigger genomes (Supplementary Fig. 3c); this result suggested ecological specificity for these viruses. For the VFGs, the most frequent VFGs were enzymes involving lipopolysaccharide (LPS) synthesis (VF0124) and PblA (VF1089, a Streptococcal phage-encoded protein that mediates binding to human platelets in the pathogenesis of infective endocarditis^34^) (Fig. 3g). In particular, LPS synthesis factor VF0124 was mainly encoded by the Podoviridae members, suggesting high pathogenicity of this viral family.

### Gut virome profiling and sex- and age-related variations

To demonstrate the utility of the cnGVC in quantitative analyses of the gut viral community, we profiled the composition of 67,096 vOTUs in all 11,327 bulk and viral metagenomic samples based on reads mapping. On average, 19.8% (interquartile range [IQR] = 17.7-22.0%) of bulk-metagenome reads could be recruited by cnGVC, which was significantly higher than any other gut viral catalogues (Fig. 4a-b; Supplementary Fig. 6a). In viral metagenomic samples, cnGVC recruited average 56.7% (IQR = 17.8-92.9%) of reads, which increased substantially compared with the existing catalogues (Fig. 4b; Supplementary Fig. 6b). For both bulk and viral samples, a large proportion of viral relative abundances (an average of 63.4% for bulk and 59.3% for viral samples) were composed by family-level unclassified viruses. Viruses belonging to Siphoviridae and Myoviridae were the most dominated members in bulk metagenomic samples, with average relative abundances of 18.7% and 8.9%, respectively, while the following Quimbyviridae and crAss-like viruses presented average abundances of 4.5% and 1.9% respectively (Fig. 4a). In viral metagenomic samples, viruses belonged to Microviridae (average relative abundance, 17.1%) were the most dominant members, followed by Myoviridae (9.6%), Siphoviridae (7.7%), and crAss-like (2.6%) viruses (Fig. 4b). The substantial higher level of Microviridae viruses in the viral samples may because that the VLP technology is more inclined to target free viral particles, agreeing with the previous studies^6, 35^.

**Fig. 4.**
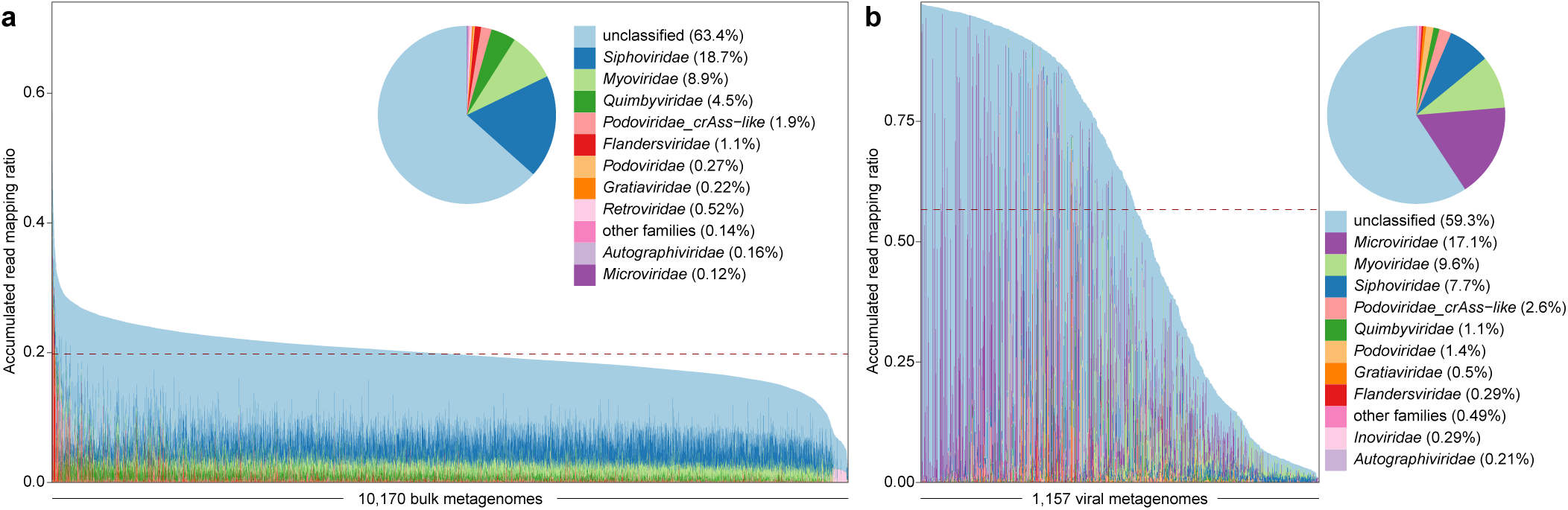
Proportion of metagenomic reads mapped into the cnGVC. **a-b,** Bar plot showing the accumulated read mapping ratio of bulk metagenomes **(a)** and viral metagenomes **(b)** used in this study. Inset pie plots show the overall read proportions at the viral family level.

Benefiting from the large datasets, we sought to explore whether the sex and age of host could impact the gut virome. This analysis was performed based on the viromes from bulk metagenomes of a total of 4,278 healthy subjects or subjects with low-risk diseases (see later for the definition) to void the effects of high-risk diseases (Supplementary Fig. 2b). Principal coordinates analysis (PCoA) based on the Bray-Curtis distance of the vOTU-level community composition revealed that both host sex and age have a modest but significant contribution to the gut virome (Fig. 5a-b). Likewise, permutational multivariate analysis of variance (PERMANOVA) showed that sex and age explained 0.28% (adonis p < 0.001) and 0.86% (adonis p < 0.001) of the overall virome variability, respectively (Fig. 5c). And these effect sizes were still significant after adjusting for study heterogeneity, host location, and body mass index (BMI).

**Fig. 5.**
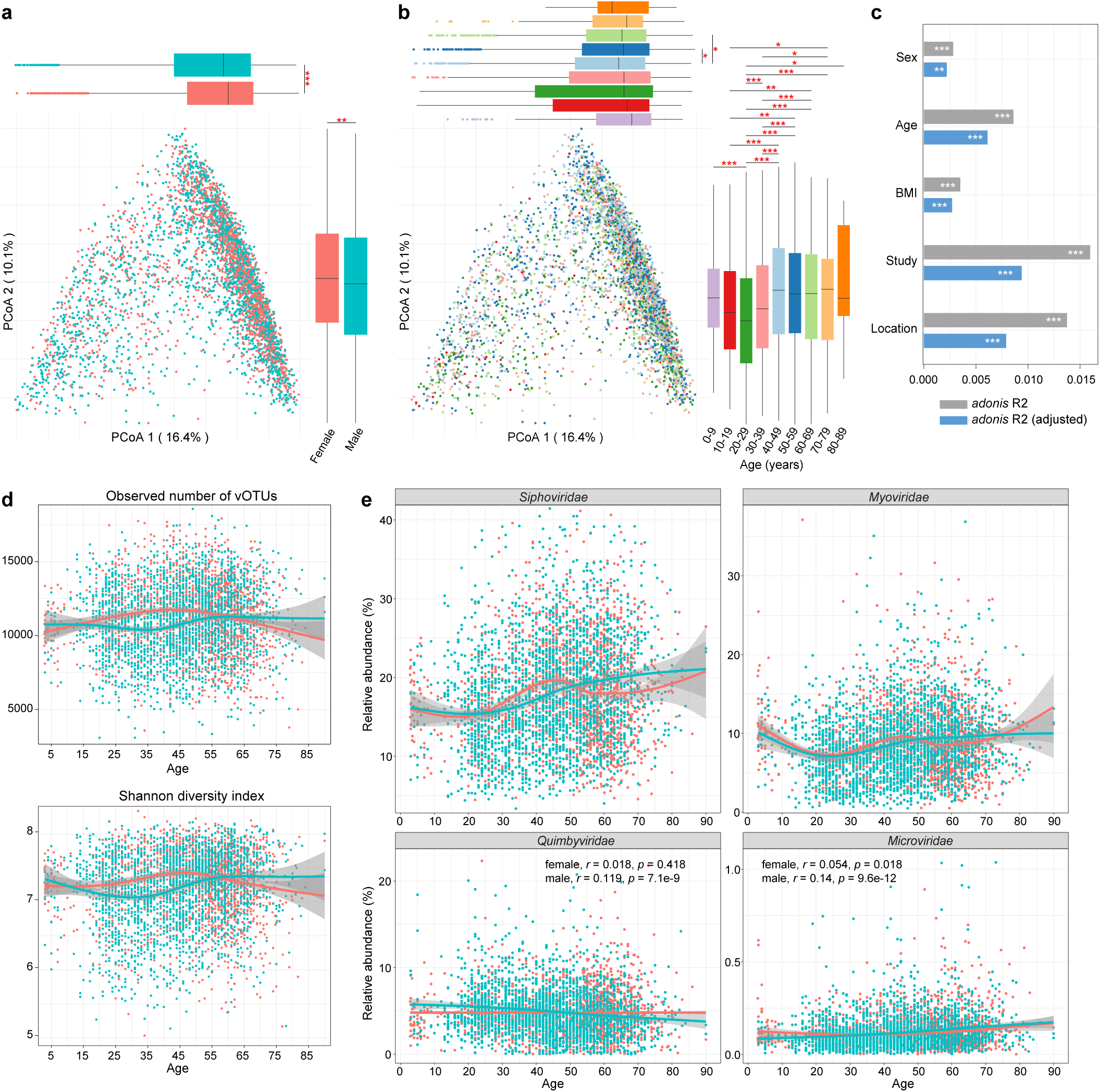
Sex- and age-related variations of the gut virome. **a-b,** Principal coordinates analysis of the gut viromes of 4,278 healthy subjects or subjects with low-risk diseases grouped by their sex **(a)** and age stages **(b)**. Samples are shown at the first and second principal coordinates (PC1 and PC2), and the ratio of variance contributed by these two PCs is shown. The below and left boxplots show the sample scores in PC1 and PC2 (boxes show medians/quartiles; error bars extend to the most extreme values within 1.5 interquartile ranges). Wilcoxon rank-sum test: *, *p*<0.05; **, *p*<0.01; ***, *p*<0.001. **c,** Permutational multivariate analysis of variance showing the effect size of sex, age, and other confounding factors on the gut virome of all investigated samples. For each factor, the raw effect size (*adonis* R^2^) and the effect size after adjusting for other factors (adjusted *adonis* R^2^) are shown. *Adonis* analysis with 1,000 permutations: *, *p*<0.05; **, *p*<0.01; ***, *p*<0.001. **d,** Scatter plots showing the sex-related trajectories of gut virome richness (upper panel) and diversity (bottom panel) at different ages. **e,** Scatter plots showing the sex-related trajectories of the top 4 dominant viral families at different ages. For **d** and **e**, points indicate samples grouped by females (red) and males (green), and smooth curves are formed based on the diversity indexes and the ages of the samples using the *geom_smooth* function in the R platform.

Using the vOTU profiles, we evaluated the gut viral richness (estimated by the observed number of vOTUs) and diversity (Shannon index) of these subjects across different age stages. Overall, the females exhibited a higher viral richness and diversity than the males (Wilcoxon rank-sum test p << 0.001 for two indexes), and this difference was mainly reflected at the age stages of 30-39 and 40-49 years (Fig. 5d; Supplementary Fig. 7). Across the life stages, we found that the gut viral richness and diversity of females are mainly increase under 40 years old (richness, r = 0.158, p = 4.2×10^-4^; diversity, r = 0.113, p = 0.0095) and dropped above 40 (richness, r = 0.136, p = 2.2×10^-7^; diversity, r = 0.115, p = 1.5×10^-5^) (Supplementary Fig. 8a). However, for males, their gut viral richness and diversity were relatively stable under 30 years old, but likely increase above 30 (richness, r = 0.128, p = 2.9×10^-8^; diversity, r = 0.172, p = 8.6×10^-14^; Supplementary Fig. 8b).

Sex- and age-related trajectories of the gut virome at the family level also showed some meaningful trends. For example, in both female and male individuals, we found that the two most dominant families, Siphoviridae and Myoviridae, have the lowest abundances at approximately 20 years old, while their abundances were relatively higher in middle-aged and elderly individuals (Fig. 5e). Siphoviridae was significantly enriched in the viromes of females at the age stages of 30-39 and 40-49 years compared with those of males (Wilcoxon rank-sum test q < 0.05), and were more deficient in females at 50-59 and 60-69 years (q < 0.05; Supplementary Fig. 9). Myoviridae was more abundant in females at 30-39 and 40-49 years than in males, while it was deficient in females at 50-59. For other virus types, Quimbyviridae viruses were decrease across ages in males but not in females, whereas Microviridae viruses were likely to increase across ages in both females and males (Fig. 5e).

### Diversity and compositional patterns of the gut virome across common diseases

Having catalogued and profiled the gut virome, we next wanted to explore the associations between gut virome and common diseases based on 36 case-control studies (Supplementary Fig. 2b). For each surveyed study, the samples were filtered under exclusion criteria such as 1) nonstandard disease definitions, 2) abnormal BMI (for samples with available phenotypic data), and 3) low metagenomic data amount or low extreme virus proportion (see Methods), which resulted in a total of 6,314 fecal samples spanning 28 disease or unhealthy statuses (40 case-control comparisons) for follow-up analyses (Supplementary Table 1). Among these diseases, cardiometabolic (7 diseases from 10 studies) and immune (8 diseases from 9 studies) disorders are the most popular, followed by digestive (3 diseases from 4 studies), infectious (3 diseases from 3 studies), and psychiatric (2 diseases from 3 studies) disorders and cancers (2 diseases from 4 studies).

Within-sample diversity analysis showed that the viral richness significantly decreased in the patient groups of 16 of 40 case-control comparisons, and similar, the viral diversity significantly decreased in patients of 13 of 40 case-control comparisons (Fig. 6a-b). Patients with Crohn’s disease (CD), pulmonary tuberculosis (PT), COVID-19 infection, and several immune (i.e., ankylosing spondylitis [AS], Graves’ disease [GD], gout, and systemic lupus erythematosus [SLE]) or cardiometabolic (i.e., hypertension, metabolic unhealthy obesity [MUO]) diseases exhibited a decrease in both viral richness and diversity. Conversely, only 3 of 40 case-control comparisons which spanned two diseases, atrial fibrillation (AF) and Parkinson’s disease, had a significant increase in viral richness or diversity.

**Fig. 6.**
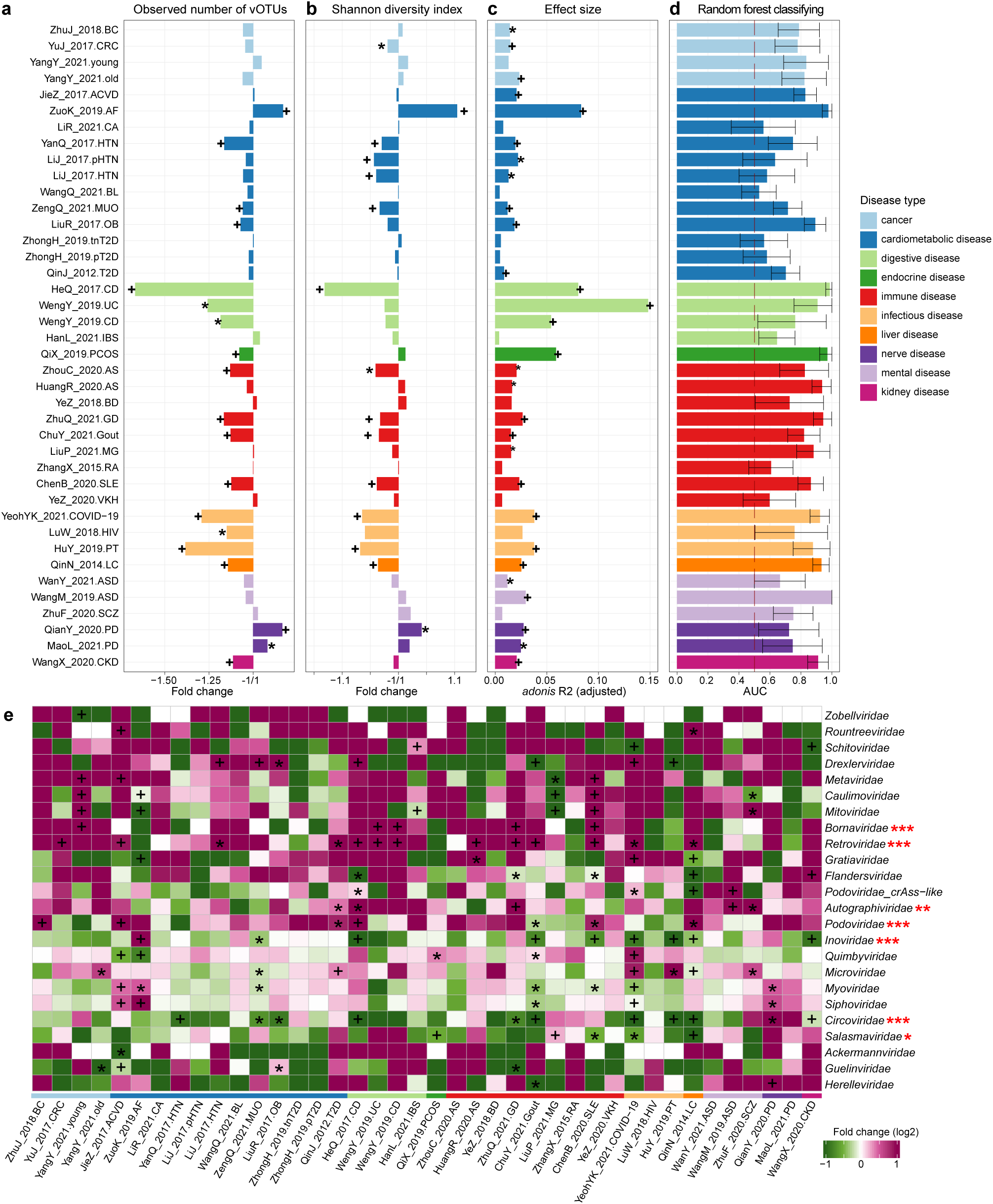
Alterations of the gut virome across common diseases. **a-d,** Bar plot showing the fold changes of gut virome richness **(a)** and diversity **(b)**, the disease-related effect size **(c)**, and the within-study AUCs **(d)** of 40 case-control comparisons. Diseases are colored by the disease types. For **a** and **b**, Wilcoxon rank-sum test: *, *p*<0.05; +, p<0.01. For **c**, *adonis* analysis with 1,000 permutations: *, *p*<0.05; +, *p*<0.01. For **d**, the dashed line shows an AUC of 0.50, and the error bars show the 95% confidence interval of the AUC values. **e,** Heatmap showing the fold changes of each viral family within 40 case-control comparisons. Fold change >0, enriched in cases; fold change <0, enriched in controls. Wilcoxon rank-sum test: *, q<0.05; +, q<0.01. The disease types of each case-control comparison are shown by bottom colors (legend following **a-d**). The red asterisks on the right represent the meta-analysis results of viral families (see Supplementary Fig. 10 for detail): *, *q*<0.05; **, *q*<0.01; ***, *q*<0.001.

PERMANOVA analysis showed that 29 of 40 case-control comparisons, spanning 22 disease or unhealthy statuses have significantly altered the overall structure of the gut virome (adonis p<0.05) (Fig. 6c). Patients with IBD showed the greatest variations in the gut virome, with effect sizes of 14.8%, 8.0%, and 5.4% in the studies WengY_2019.UC (p<0.001), HeQ_2017.CD (p<0.001), and WengY_2019.CD (p<0.001). Furthermore, we carried out random forest classifying to distinguish cases and controls within each study based on their gut viral profiles. The classifiers achieved a high discriminatory ability (AUC>0.80) in 19 of 40 case-control comparisons and a moderate discriminatory ability (AUC = 0.70-0.80) in the remaining 11 case-control comparisons (Fig. 6d). These findings demonstrated profound changes in the gut viral composition across many diseases that substantially differ in clinical manifestations and pathogenesis.

On the other hand, combining the diversity, PERMANOVA, and random forest classifying results, we found that 7 diseases, including carotid atherosclerosis (CA), bone mass loss (BL), irritable bowel syndrome (IBS), Behcet’s disease (BD), rheumatoid arthritis (RA), Vogt-Koyanagi-Harada disease (VKH), and schizophrenia, do not have a considerable change in both the gut viral diversity and composition; these diseases are likely low-risk diseases in the gut virome scope.

To further uncover the gut viral signatures, we next compared the gut viromes between patients and controls for each disease at the family level. Using the Wilcoxon rank-sum test, we found that the Siphoviridae viruses are significantly enriched in patients with atherosclerotic cardiovascular disease, AF, and Parkinson’s disease (q < 0.05) compared with their corresponding healthy controls and reduced in patients with gout and COVID-19 infection (q < 0.05; Fig. 6e). Myoviridae were very similar to Siphoviridae in viral signatures, with additional reductions in the patients of MUO and SLE (q < 0.05). Microviridae were remarkably proliferated in patients with old CRC, type 2 diabetes (T2D), COVID-19 infection, PT, and schizophrenia but only reduced in MUO subjects. Interestingly, several viral families showed the same tendency in most diseases. For example, Retroviridae was significantly enriched in the patients from 13 case-control comparisons that spanned 11 diseases, Autographiviridae and Bornaviridae were enriched in the patients of 5 different diseases, and Podoviridae was enriched in 6 diseases and depleted in 1. Random effects meta-analysis supported that these 4 families are significantly more abundant in patients across all diseases (q < 0.05; Supplementary Figure 10). Likewise, Circoviridae, Inoviridae, and Salasmaviridae were decreased in a diverse range of disease patients, with an overall deficiency across all diseases (meta-analysis q < 0.05). Collectively, these findings suggested the existence of shared viral signatures for health.

### Universal viral signatures of common diseases

Given that most of the investigated diseases exhibited significant alterations in the overall viral communities and in certain viral families, therefore, we wanted to investigate the universal viral signatures of these diseases at the vOTU level. A total of 7,161 vOTUs that differed in relative abundances across 36 case-control studies were identified based on the combination of meta-analysis and direct comparison between all case and control subjects (random effects meta-analysis q < 0.01 and cases vs. controls q < 0.01; Fig. 7a; Supplementary Table 3). 2,531 of these differential vOTUs were more abundant in subjects with diverse diseases, while 4,630 of them were enriched in healthy individuals. Both disease-enriched and control-enriched vOTUs were mainly composed of Siphoviridae, Myoviridae, and unclassified members. Consistent with the aforementioned findings at the family level, members of Podoviridae (disease-enriched vs. control-enriched vOTUs, 131 vs. 3) and Retroviridae (126 vs. 0) were frequently enriched in disease subjects, and members of Inoviridae (2 vs. 48) mainly appeared in the control-enriched vOTUs (Fig. 7b; Supplementary Fig. 11a). Besides, 40 vOTUs of Quimbyviridae were disease-enriched but only 11 of them were control-enriched (Fisher’s exact test q = 9.5×10^-10^), and 13 vOTUs crAss-like were control-enriched but none crAss-like member was disease-enriched (q = 0.007). When assigning the disease-differential vOTUs into prokaryotic hosts, we found that the control-enriched vOTUs had a large proportion of members that were predicted to infect Ruminococcaceae (17.6% of 4,630 control-enriched vOTUs) and Oscillospiraceae (10.3%), whereas only 0.6% and 1.5% of disease-enriched vOTUs were members of these two families, respectively (Fig. 7b; Supplementary Fig. 11b). Notably, a large proportion of Ruminococcaceae-hosted vOTUs was predicted to infect bacteria of the Faecalibacterium genus (n = 413 vOTUs; Supplementary Fig. 11c), a well-known SCFA-producer taxon that showed beneficial effects in multiple common disorders^36, 37^; suggesting a potential link between Faecalibacterium phages and health. Conversely, the disease-enriched vOTUs contained a considerably high proportion of viruses that were predicted to infect Enterobacteriaceae (n = 344, corresponding to 13.6% of 2,531 disease-enriched vOTUs), Streptococcaceae (2.2%), Fusobacteriaceae (1.3%), Erysipelotrichaceae (1.3%), and Erysipelatoclostridiaceae (1.0%), while phages of these bacteria had rarely appeared in the control-enriched vOTUs. At the genus level, the phages of Escherichia were the most frequent in disease-enriched vOTUs (Supplementary Fig. 11c).

**Fig. 7.**
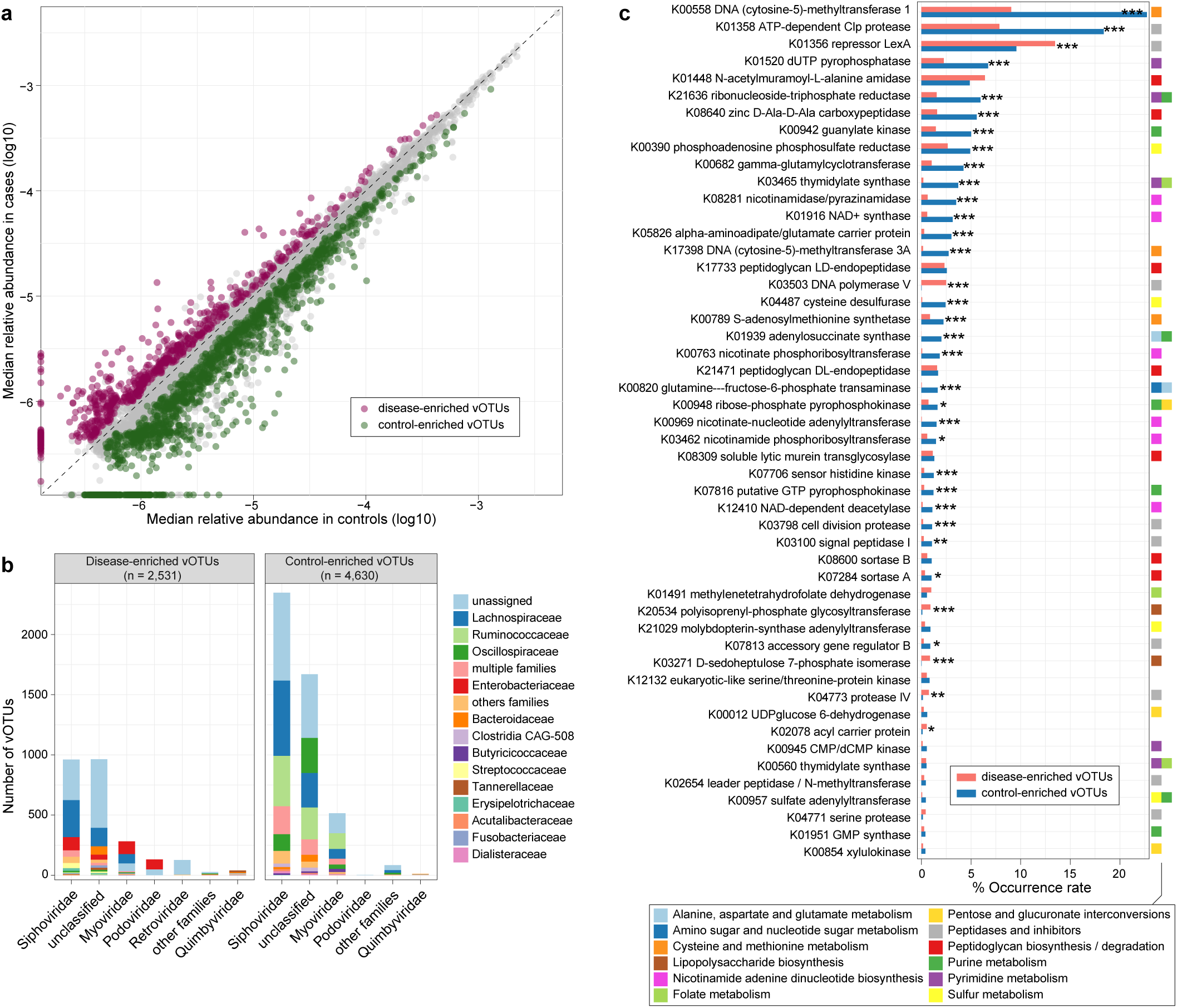
The disease-associated viral signatures. **a,** Scatter plot of median relative abundances of vOTUs in all investigated disease and control individuals. Grey points represent vOTUs not differentially abundant between two groups, and red and green points represent differentially abundant vOTUs. **b,** Distribution of taxonomic annotation and host assignment of the disease-enriched and control-enriched vOTUs. The vOTUs are grouped at the family level, and the prokaryotic host taxa are also shown at the family level. The number of vOTUs that had more than one predicted host is labeled by red color. **c,** Occurrence rate of 50 most frequent AMGs in all disease-associated vOTUs. The functional categories of AMGs are shown by colored squares. Statistical test was performed using Fisher’s exact test: *, *q*<0.05; **, *q*<0.01; ***, *q*<0.001.

To elucidate the functional and metabolic capabilities of the gut viral signatures of common diseases, we compared the profiles of AMGs between disease-enriched and control-enriched vOTUs at the enzyme level (Supplementary Table 4). The preliminary comparison showed that the control-enriched vOTUs encoded a higher frequency of AMGs than the disease-enriched vOTUs (Supplementary Fig. 12a), suggesting that they may involve in the metabolism of more substances in the human gut. In particular, 34 of the 50 most frequent AMGs differed in frequency between disease-enriched and control-enriched vOTUs (Fig. 7c). The control-enriched vOTUs had a higher frequency of enzymes involving biosynthesis of nicotinamide adenine dinucleotide (NAD+) (n = 6 enzymes, Supplementary Fig. 12b), cytosine/methionine metabolism (DNA cytosine-5-methyltransferase K00558/K17398 and S-adenosylmethionine synthetase K00789), folate metabolism (thymidylate synthase K03465/K00560 and methylenetetrahydrofolate dehydrogenase K01491), and assimilatory sulfate reduction (phosphoadenosine phosphosulfate reductase K00390 and sulfate adenylyltransferase K00957) than the disease-enriched vOTUs, whereas the two enzymes involving to LPS biosynthesis, polyisoprenyl-phosphate glycosyltransferase K20534 and D-sedoheptulose 7-phosphate isomerase K03271, were more frequent in the disease-enriched vOTUs. A higher level of viral-encoded NAD+ synthesis capacity was also observed in the gut virome of patients with CKD (unpublished data), probably linked to their roles in phage DNA replication and exploitation of the host metabolic pathways and biochemical processes during viral infection^38^, which needs to be validated by subsequent studies.

### Gut viral signatures as a predictor for health status

Finally, we tested the ability of gut viral signatures in predicting human health status. A random forest classifier was trained based on the abundances of 7,161 universal viral signatures in all 6,314 samples across 36 studies and tested using the tenfold cross-validation approach. This classifier obtained a performance of the area under the receiver operating characteristic curve (AUC) score of 0.708 (95% confidence interval [CI], 0.695-0.721) in classifying all case and control samples (Fig. 8a). In addition, it achieved an AUC of 0.731 (95% CI, 0.716-0.745) in distinguishing the patients with high-risk diseases from their corresponding control subjects, highlighting the high predictability of these patients by gut virome. Interestingly, a new classifier trained by a subset of 70 most important vOTUs generated the discriminatory power of AUC 0.698 (95% CI, 0.685-0.711) for classifying all samples and 0.718 (0.703-0.732) for high-risk diseases versus controls (Supplementary Fig. 13); this a minimal set of gut viral signatures may be used as a more feasible indicator for health.

**Fig. 8.**
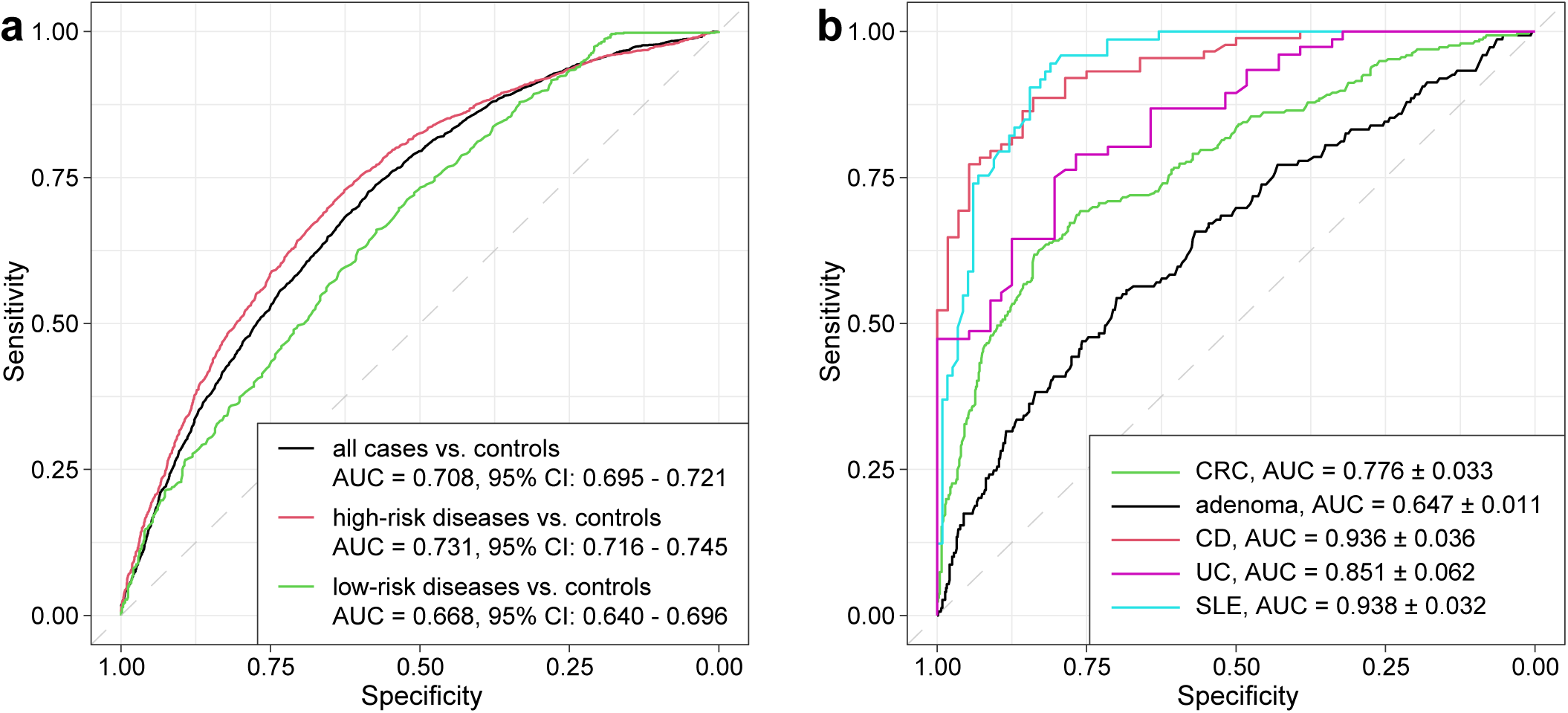
Prediction of health status using the viral signatures. **a,** Receiver operating characteristic (ROC) analysis of the classification of case/control status using the random forest model trained by 7,161 universal viral signatures. **b,** ROC analysis of the classification of case/control status in independent cohorts. The classification performance of the model was assessed by the area under the ROC curve (AUC). The AUC values and 95% confidence intervals (CIs) are shown.

We curated fecal metagenomes from three independent cohorts, including the CRC cohort (comprising 480 CRC patients, 182 adenoma patients, and 492 healthy controls from 8 studies from European, USA, and Japan populations)^39^, the IBD cohort (88 CD patients,76 UC patients, and 56 controls from the USA)^40^, and a newly-recruited cohort comprising 73 SLE patients and 116 controls from China (Supplementary Table 5), to validate the reliability of gut viral signatures. Using these large cohorts, we quantified the relative abundances of 7,161 disease-associated vOTUs and compared them between cases and controls in new cohorts. The results showed that most vOTUs had a consistent trend in mean abundance between patients and controls within each disease compared with the observation in the original datasets. For example, in the CD patients vs. controls, 95.5% (6,837/7,161) of vOTUs were more abundant in patients or controls, consistent with the original datasets, and 67.1% (4,808/7,161) of vOTUs were significantly enriched (Supplementary Fig. 14). Moreover, the discriminatory powers of the original random forest classifier on these new cohorts were 0.776, 0.647, 0.936, 0.851, and 0.938 for CRC, adenoma, CD, UC, and SLE patients versus controls, respectively (Fig. 8b). These findings suggested that the generalized disease-associated gut viral signatures identified in this study can accurately classify multiple diseases from health, even in subjects from different nations.

## Discussion

Viruses are in number the largest component of the human gut microbial community, but much about their genome, function, and role in certain diseases is unknown^41^. In this study, we reported the cnGVC which contained over 67,000 nonredundant viral sequences with high completeness (approximately 60% of viruses are ≥90% completion) and representativeness. This catalogue is, to our knowledge, the current largest viral genome catalogue for fecal metagenomes from a single population. In particular, it is even 71.3% and 55.8% larger than GPD and MGV, respectively, although the latter databases are from large-scale populations in multiple countries. Noticeably, a comparison of different gut virus catalogues revealed that more than 70% of viruses in cnGVC were newly found. This is not only due to technical differences (e.g., inclusion criteria, use of VLP datasets), but also highlights the specificity of the gut virome in the Chinese population. Overall, the construction of cnGVC demonstrated the great research value of viral identification targeting a single population.

We constructed a gene catalogue from cnGVC, which represented nearly 1.6 million non-redundant viral genes. The majority (87.4%) of the viral genes were not found in the gut prokaryotic genomes, this finding agreed with the previous observations in other habitats^42–44^ and further underscored the high functional specificity of the virome in context of the future holistic microbiome research. In addition, we identified numerous AMGs and CAZymes and a certain number of VFGs and ARGs from the viral gene catalogue. These resources may also facilitate future studies to comprehensively decipher some important viral functions such as complex polysaccharide degradation and antibiotic resistance.

Using massive fecal metagenome datasets, we found that both individuals’ sex and age have a considerably impact on the gut virome and uncovered these sex- and age-related alteration patterns. A previous study had revealed a decrease in gut viral diversity in elderly individuals compared with that in adults^6^, while this phenomenon was not evident in our data. We found that both the gut viral diversity and the compositions of dominant viral families (i.e., Siphoviridae and Myoviridae) of females have great changes at the age of 40-49, these findings are similar to the observations in the gut bacteriome and may be related to the dramatic physiological changes (e.g., changes in sex hormones or metabolism) for women after menopause^45, 46^. Based on our results, we speculated that the gut virome may play potential roles in human physiology, immunity, or metabolism; this notion is worth for future systematic investigation.

The gut virome has been implicated in a variety of human diseases, however, integrating and comparing viral signatures across diseases remains a challenge. Based on our large-scale datasets, we found that the viral richness and diversity have frequently been reduced in a variety of diseases, but only increased in a few of them. Loss of gut viral diversity was also reported in other single disease studies, such as T2D^47^ and liver disease^12^. Similarly, diversity reduction is yet a typical phenomenon in the gut bacteriome under disease status variations, which may be related to low resilience and possible dysbiosis within the microbiological ecosystem^48, 49^. Moreover, we revealed that a majority of the investigated diseases are associated with significant alterations in overall viral communities and their degree of virome alteration is diverse. These findings 1) agreed with previous studies showing substantial changes of the virome in IBD, CRC, and several other diseases, 2) strengthened the previous connections between gut virome and immune/cardiometabolic diseases, and 3) proposed several previously unexplored diseases which are alterable in virome, such as Parkinson’s disease, autism spectrum disorder, and polycystic ovarian syndrome (Figure 6a-d). Follow-up exploration of our results may generate testable hypotheses to guide future disease-specific research on its etiologies and therapeutic strategies.

Meta-analysis across all investigated diseases identified extensive gut viral signatures at both family and vOTU levels. The most significant disease-associated viral families were Podoviridae and Retroviridae, which were significantly enriched in the gut of patients with 5 and 11 different diseases, respectively, with meta-analysis q < 1×10^-4^ for both. Members of Podoviridae are mostly temperate bacteriophages, but they also encode the majority of virulence factors involving LPS synthesis as found in this study. Almost all gut Retroviruses were newly discovered by cnGVC, and their functions are still under exploration. Other universal viral signatures included a large number of disease-enriched vOTUs that were predicted to infect Enterobacteriaceae, Streptococcaceae, Fusobacteriaceae, Erysipelotrichaceae, and Erysipelatoclostridiaceae. Pathogenic Enterobacteriaceae bacteria have been confirmed as disease-common opportunistic pathogens^50, 51^, and their phages may interact with host bacteria to act on diseases. The other bacterial taxa were only observed with pathogenicity in certain specific diseases (e.g., Streptococcaceae for liver cirrhosis and Fusobacteriaceae for CRC)^52, 53^. Hence, their phages may be partially independent of bacteria to play roles in human diseases -- further research is needed to uncover the exact role of these viral clades in human diseases. Moreover, functional analyses revealed that some viral functions, such as NAD+ synthesis, have been widely associated with diseases, highlighting the value of viral functions in common diseases. In addition, we found that the gut viral signatures have high predictive power for disease status across all samples, and this predicting performance is comparable with recent bacterial level studies^54^. On the test datasets, we also found that these signatures have high reproducibility in multiple external populations. Collectively, the broad and universal viral signatures obtained from our study may be useful for future disease mechanism, intervention, and phage therapy efforts.

## Methods

### Human fecal metagenomic datasets and public databases

We reviewed a series of studies based on human fecal metagenomic samples by searching for given keywords (e.g., “gut metagenome”, “fecal metagenome”, “stool metagenome”, “viral-like particle”, “VLP metagenome”) in the PubMed search system, and manually selected out 50 studies of Chinese cohorts with publicly available metagenomic data until Marth 2022. These studies provided 11,327 fecal metagenomic samples containing over 95 Tbp of high-throughput sequencing data. Detailed information of 50 studies was shown in Supplementary Table 1.

The five public databases of human gut viral and microbial genomes were involved in this study: 1) the Gut Virome Database (GVD) downloaded from https://datacommons.cyverse.org/browse/iplant/home/shared/iVirus/Gregory_and_Zablocki_GVD_Jul2020/GVD_Viral_Populations6; 2) the Gut Phage Database (GPD) downloaded from http://ftp.ebi.ac.uk/pub/databases/metagenomics/genome_sets/gut_phage_database/1; 3) the Metagenomic Gut Virus (MGV) catalogue downloaded from https://portal.nersc.gov/MGV/20; 4) the Cenote Human Virome Database (CHVD) downloaded from https://zenodo.org/record/449888455; and 5) the Unified Human Gastrointestinal Genome (UHGG) collection downloaded from http://ftp.ebi.ac.uk/pub/databases/metagenomics/mgnify_genomes/human-gut/v1.0/uhgg_catalogue/56.

### Sequence preprocessing and metagenome assembly

Raw reads were quality filtered and trimmed using fastp (v0.20.1)^57^ with the parameters ‘-l 60 -q 20 -u 30 -y –trim_poly_g’ (for samples with read length ≤ 100 bp) or ‘-l 90 -q 20 -u 30 -y –trim_poly_g’ (for samples with read length > 100 bp), and human contamination was removed by mapping quality-filtered reads with bowtie2 (v2.4.1)^58^ against the human reference genome (GRCh38). The remaining clean reads in each sample were assembled into contigs using MEGAHIT v1.2.9^59^ with the different lists of k-mer values that were set based on read length.

### Identification and decontamination of viral sequences

We performed an integrated homology- and feature-based pipeline to identify viral sequences based on our previously developed methodologies ^7, 39, 60, 61^. In brief, we first removed assembled contigs whose prokaryotic genes were over half of their genes and ten times greater than the number of viral genes based on CheckV (v0.7.0)^22^ assessment. The remaining assembled contigs were used for viral identification when they were deemed to meet any of the following criteria: 1) contig had at least one viral gene, and its viral genes were greater than the number of prokaryotic genes based on CheckV (v0.7.0) assessment; 2) contig had a DeepVirFinder (v1.0)^62^ score of > 0.90 and p-value of < 0.01; 3) contig was identified as viral sequence by VIBRANT (v1.2.1)^30^ with default options. After screening, the contigs were identified as potential viral sequences. In addition, given the limited value of viral sequences with low completeness to subsequent works, we removed the viral sequences with CheckV completeness of <50%. According to the previous study^6^, we further performed a decontamination process for the remaining viral sequences based on the ratio of bacterial universal single-copy orthologs (BUSCO ratio)^63^. The hmmsearch program^64^ was used to search BUSCO genes within each viral sequence with default parameters, and the BUSCO ratio of the viral sequence was calculated as the number of BUSCO divided by the total number of its genes. Viral sequences with >5% BUSCO ratio were then removed. Finally, totaling 426,496 viral sequences were retained as highly credible viral genomes for follow-up analysis.

### Viral clustering and gene prediction

Referring to the clustering method provided by our prior study^60^, we clustered viral sequences at 95% average nucleotide similarity threshold (≥70% coverage) and resulted in a nonredundant gut virus catalogue comprised of 67,096 vOTUs. Approximately 22.0 million putative protein sequences in all vOTUs were predicted via Prodigal (v2.6.3)^65^ with the parameter ‘-p meta’, and were clustered into nearly 1.6 million nonredundant protein sequences using MMseqs2 (v12.113e3) easy-linclust mode^66^ with the parameters ‘--min-seq-id 0.9 --cov-mode 1 -c 0.8 --kmer-per-seq 80’.

### Taxonomic classification, host assignment, and functional annotation

We ran the Diamond program (v2.0.13.151)^67^ with parameters ‘--id 30 --query-cover 50 --min-score 50 --max-target-seqs 10’ to taxonomically classify vOTUs by aligning protein sequences against an integrated viral protein database that was derived from Virus-Host DB^68^ downloaded in March 2022, crAss-like phage proteins provided by Guerin et al’s study^69^, viral proteins provided by Benler et al’s study^25^, and human viral proteins provided by Ye et al’s study^70^. A vOTU was assigned into the known viral family when over one out of five proteins were annotated to the same family.

Host assignment of vOTUs was performed based on their homology to genome sequences or CRISPR spacers of the comprehensive unified human gastrointestinal genome (UHGG) database^56^. For the homologous alignments, the viral sequence was aligned against prokaryotic genomes in UHGG using BLASTn with the parameters ‘-evalue 1e-2 -num_alignments 999999’, and assigned a host when >30% region of the viral sequence was matched to the corresponding host genome at > 90% nucleotide identity and <1e-10 e-value. For CRISPR-spacer matches, we carried out the detection of host CRISPR-spacer sequences using MinCED v0.4.2 with the parameter ‘-minNR 2’, and then assigned a host to vOTU when the host CRISPR spacer and viral sequences could be aligned by BLASTn at >45 bit-score and <1e-5 e-value.

To investigate the functional properties of viruses, we implemented the functional annotation of viral proteins based on the KEGG^71^, CAZy^72^, VFDB^73^, and combined antimicrobial resistance databases (comprised of CARD v3.1.0^74^, MEGARes v2.0^75^, ResFinder^76^, and ARG-ANNOT^77^ databases), respectively. For the KEGG and CAZy databases, the protein was assigned a functional ortholog on the basis of the best-hit gene in the database using Diamond with parameters ‘-e 1e-5 --query-cover 50 --subject-cover 50 --min-score 50’. For the VFDB, the protein was assigned a virulence factor on the basis of the best-hit gene in the database using Diamond with parameters ‘--query-cover 50 --id 60’. For the combined antimicrobial resistance database, viral proteins were aligned against the combined database using Diamond with parameter ‘-e 1e-2’. Notably, we selected different threshold criteria of protein identity for different types of resistance proteins to annotate viral proteins (>70% for multiple drug resistance proteins; >90% for beta-lactamases; >80% for the others).

### Phylogenetic analysis

We performed the genome-based phylogenetic analyses for vOTUs with CheckV completeness of > 90% using ViPTreeGen (v1.1.2)^78^ with the default parameters. The phylogenetic tree was visualized by iTOL (v6). In addition, we calculated the phylogenetic diversity (PD) of each viral family using the pd function within the R picante^79^ package based on the proteomic tree generated by ViPTreeGen.

### Taxonomic profiles

To characterize the composition of gut virome, clean reads in each metagenome were mapped to vOTUs in cnGVC via bowtie2 with the parameters ‘--end-to-end --fast --no-unal -u 10000000’. The abundance of each vOTU was calculated as the number of reads mapped to this vOTU. The relative abundance of each vOTU was calculated as its abundance divided by the number of reads mapped to all vOTUs in cnGVC. We also calculated the relative abundances of viral families in each metagenome by adding up the relative abundances of vOTUs belonging to the same family.

### Statistical analysis

Statistical analyses and data visualization were carried out via R language (v4.0.3).

#### Alpha diversity

The two alpha diversity estimates (i.e., the Shannon index and the observed number of vOTUs) were measured based on the relative abundance profiles at the vOTU level. Shannon index was estimated using the diversity function within the vegan package. The observed number of vOTUs was calculated as the number of vOTUs with relative abundance >0 in each metagenome.

#### Multivariate statistics

The Bray-Curtis distances between samples were calculated based on the relative abundance profiles at the vOTU level using the vegdist function within the vegan package. Principal coordinate analysis (PCoA) was carried out based on between-sample Bray-Curtis distances using the pcoa function within the ape package. Permutational multivariate analysis of variance (PERMANOVA) was implemented using the adonis function, and then adonis R^2^ was adjusted using the RsquareAdj function.

Correlation analyses. Correlation coefficients for metadata and viruses-associated variables were estimated using the cor.test function with the parameter ‘method = pearson’. Smooth curves were formed using the geom_smooth function with default parameters.

Statistical tests. Wilcoxon rank-sum tests were performed on differences in virome diversities and viral relative abundances between controls and patients in 50 studies via the wilcox.test function. Furthermore, we performed random effects meta-analysis to identify universal viral signatures of common diseases. In brief, we first performed the arcsine square root transformation for the relative abundances of vOTUs, and then used Hedges’ g effect sizes to assess the difference in the transformed values of each vOTU between controls and patients via the escalc function with the parameter ‘measure=SMD’. Meanwhile, study heterogeneity was quantified based on I^2^ statistic and tested by Cochran’s Q-test using the rma function within the metafor package. For comparison analyses of occurrence rates of AMGs between disease- and control-enriched vOTUs, the occurrence rate of AMG was calculated as the number of vOTUs with AMG divided by the total number of vOTUs within the corresponding group, and then statistically tested using the fisher.test function.

#### Classification models

We built the random forest model to test the ability of gut viral signatures in predicting human health status using the randomForest function, and the performance of the model was verified by repeating 10 times 10-fold cross-validation.

### Data and code availability

The viral sequences and annotation files of cnGVC have been deposited in the GitHub website with URL:

https://github.com/yexianingyue/GV_common_diseases/. The original codes used in the paper are provided in the same URL. All other data reported in this paper will be shared by the corresponding authors upon request.

## Acknowledgements

This work was supported by the National Natural Science Foundation of China (No. 82225048 and 81902037), Beijing University of Chinese Medicine (No. 5050071720001 and 2180072120049).

## Author contributions

S.L., W.S., and X.M. conceived the study. S.L. and Yu.Z., performed data collection. S.L., Yu.Z., R.G., Q.L., F.C., Jin.M., and G.W. performed the gut virome analyses. Q.Y. and P.Z. participated in development of analytical methods. Q.Y., L.C., S.F., R.L., W.Y., Ya.Z., and Jie.M. performed sample collection and experiments. S.L. drafted the manuscript. Z.L., J.L., C.C., and H.U. helped drafting the manuscript. All authors revised the article and approved the final version for publication.

## Competing interests

The authors declare no competing interests.

## Notes

### Competing Interest Statement

The authors have declared no competing interest.

