## Supplemental Figure for "Cataloguing and profiling of the gut virome in Chinese populations uncover extensive viral signatures across common diseases"

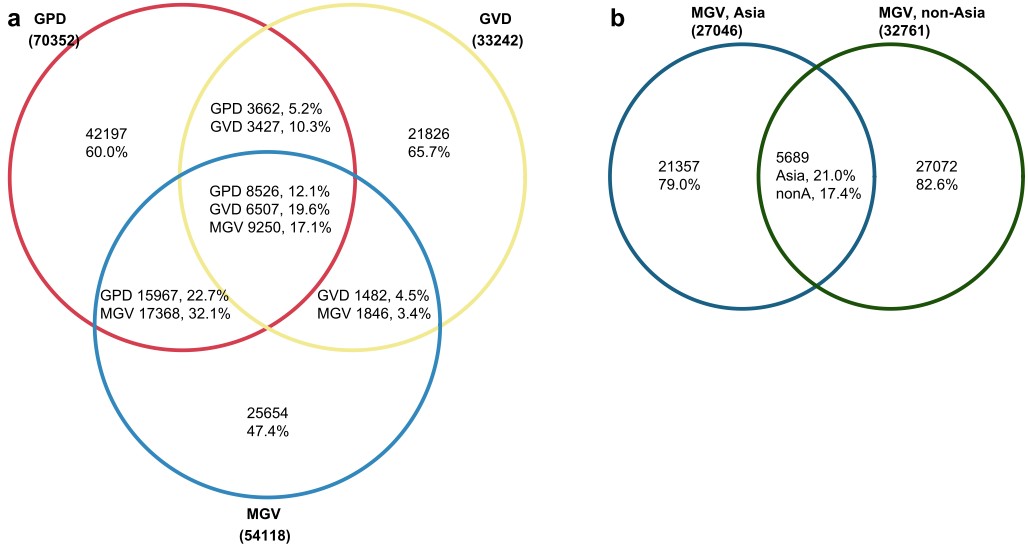

**Supplementary Fig. 1| Substantial differences of the gut viruses from different sources. a,** Venn plot showing the sharing relationship of viruses from three publicly gut virus catalogues. **b,** Venn plot showing the sharing relationship of viruses in the MGV catalogue origin from Asia and non-Asia samples. GVD, gut virome database; GPD, gut phage database; MGV, metagenomic gut virus.

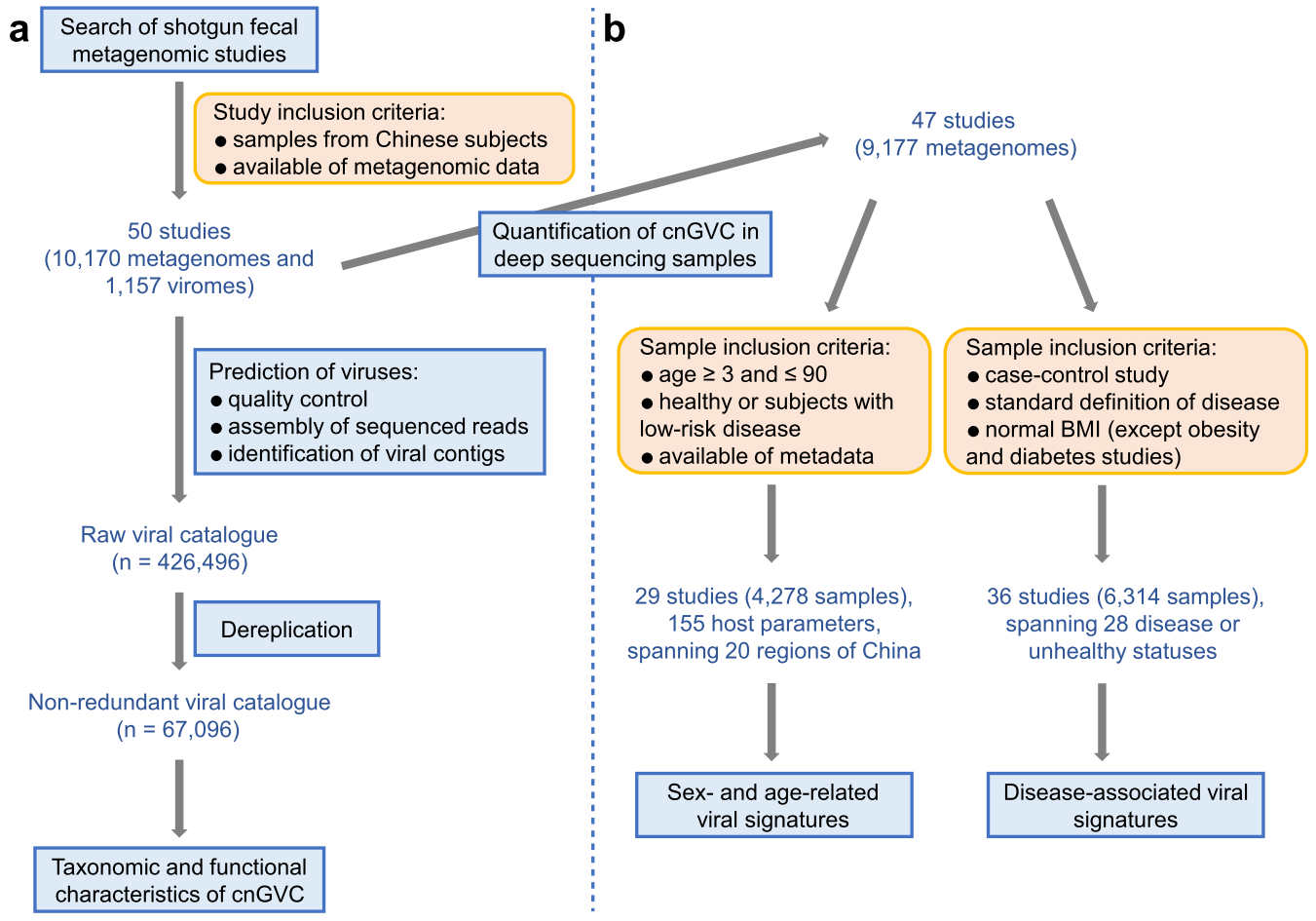

**Supplementary Fig. 2| Overview of the workflows in this study.** **a,** Workflow for the construction of the cnGVC based on 10,170 fecal bulk metagenomes and 1,157 fecal viral metagenomes deriving from 50 previously published studies. **b,** Workflow for profiling and analyzing the gut viromes among fecal bulk metagenomes.

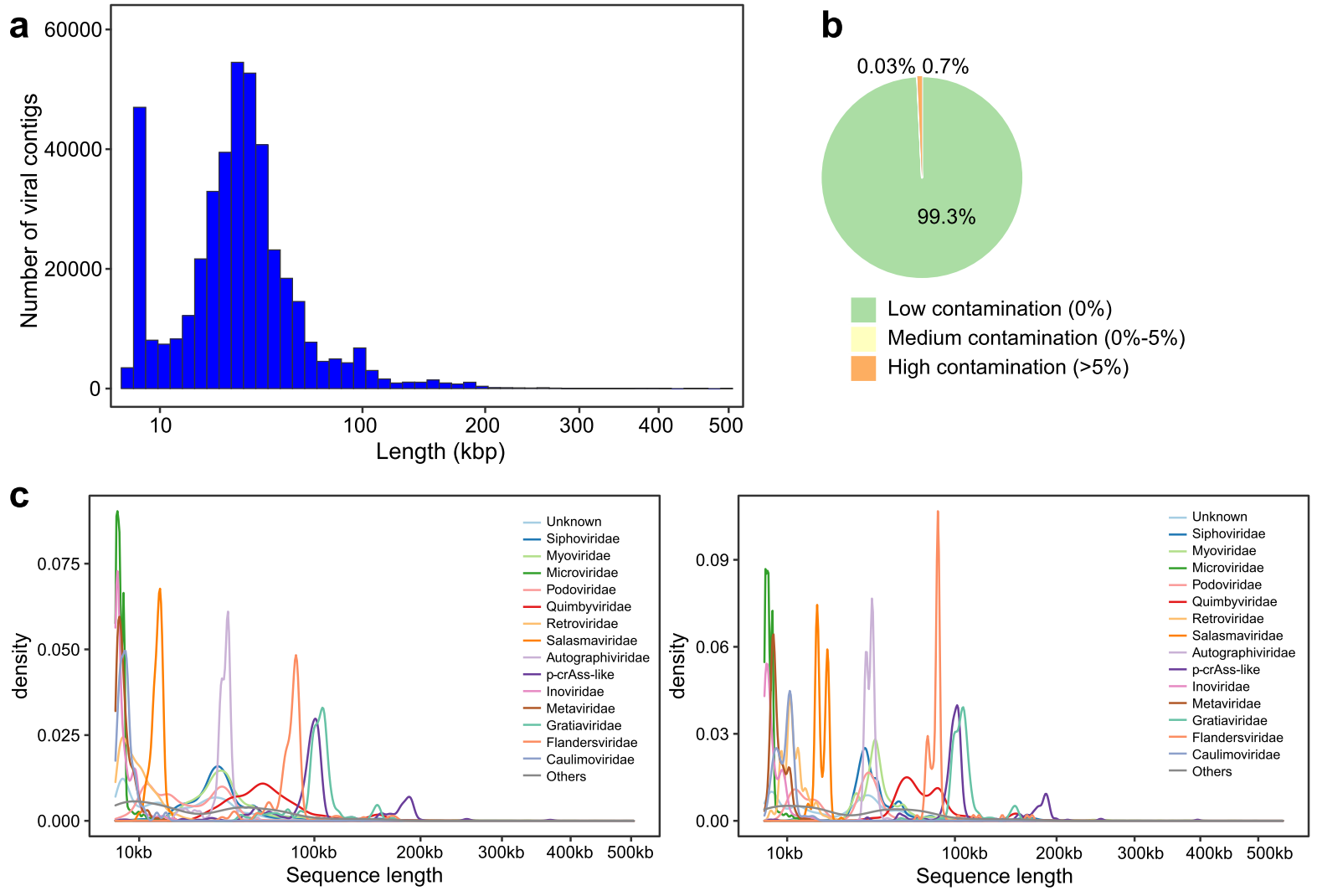

**Supplementary Fig. 3| Statistics of length and contamination of the viral genomes in cnGVC. a,** Distribution of the length of all viral genomes. **b,** CheckV-based estimation of the contamination rate of all viral genomes. **c,** Distribution of the length of viral genomes grouped at the family level. Left panel, raw viral lengths; right panel, viral lengths are adjusted by the estimated completeness.

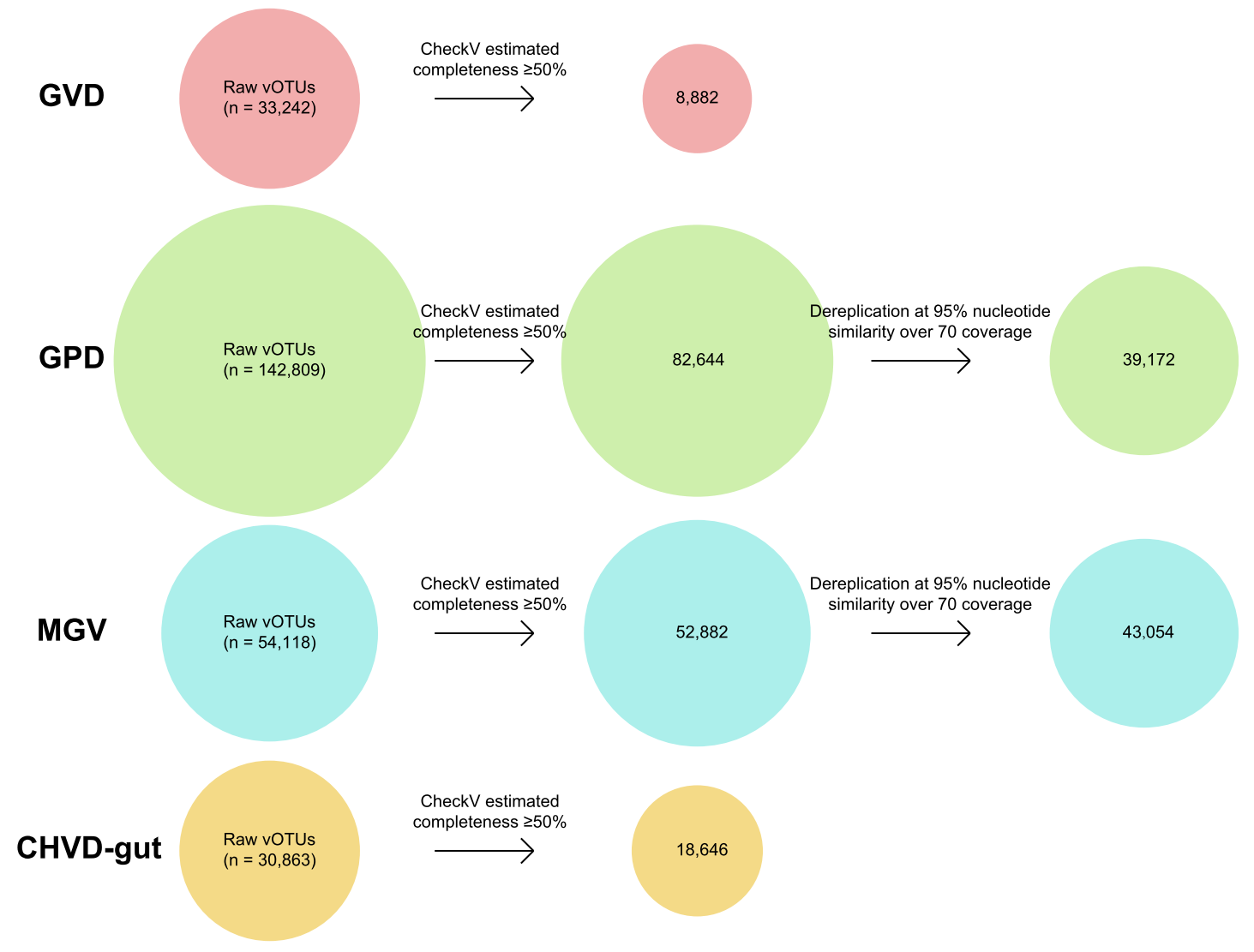

**Supplementary Fig. 4| Processing of the existing gut viral databases using a unified pipeline.** Raw vOTU sequences of each database were downloaded from the corresponding websites, and were then processed in two steps: 1) low-quality viruses with estimated completeness <50% were removed; 2) dereplicated at 95% nucleotide similarity and 70 coverage using custom scripts. The second step was not performed for GVD and CHVD-gut as they had already been dereplicated using the same parameters. The scripts for the dereplication process were available at <https://github.com/RChGO/gut_virome_db/>.

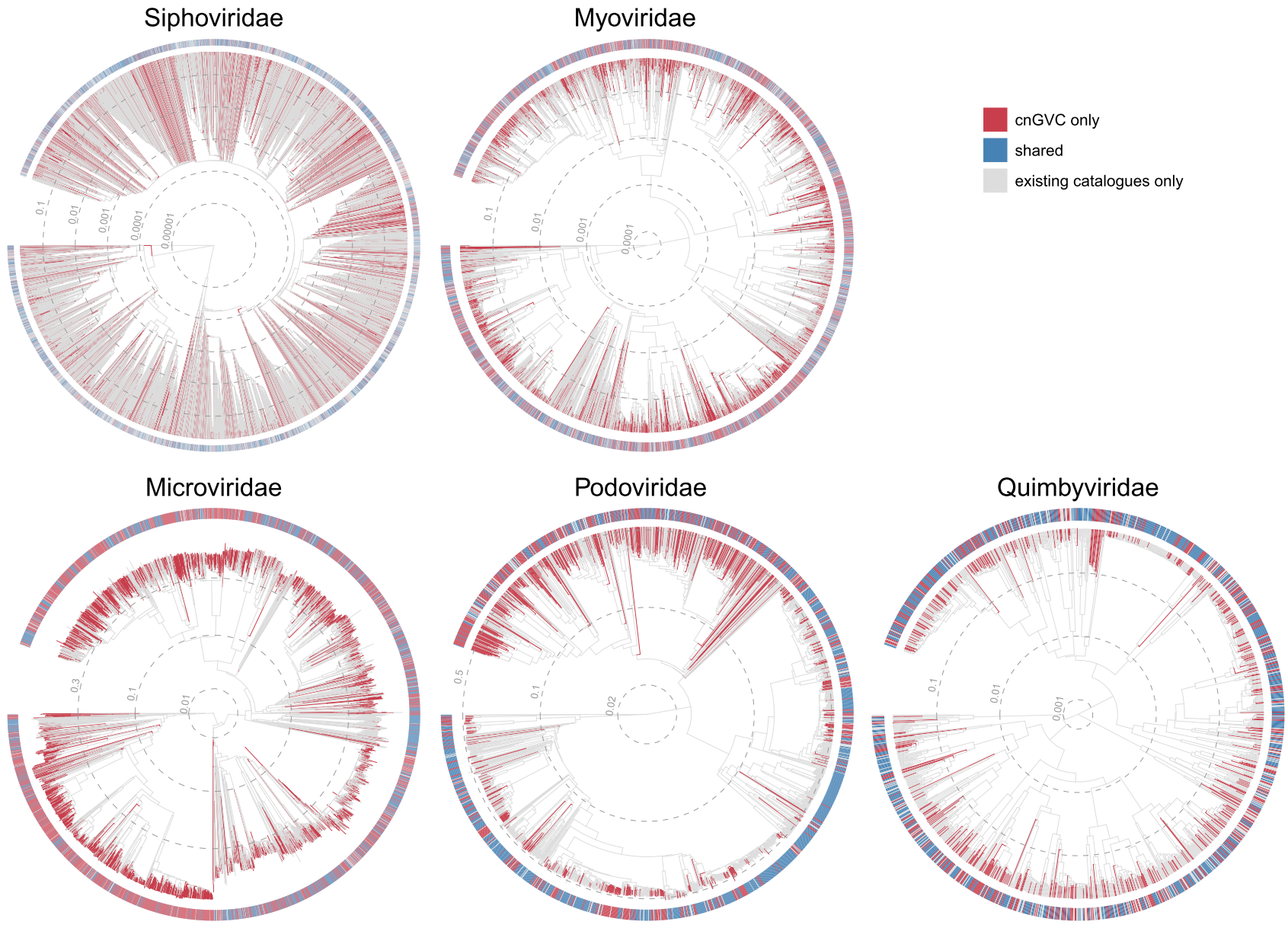

**Supplementary Fig. 5| Genome-based phylogenetic trees of 5 dominant viral families.** The tree was generated based on the protein sequences of high completeness viruses using ViPTreeGen [1]. Outer rings display the sources for each virus. For each family, we found that the newly-found vOTUs in cnGVC are broadly distributed in the major taxonomic lineages across the phylogenetic tree.

[1] Nishimura Y, Yoshida T, Kuronishi M, et al. ViPTree: the viral proteomic tree server. *Bioinformatics*, 2017, 33(15): 2379-2380.

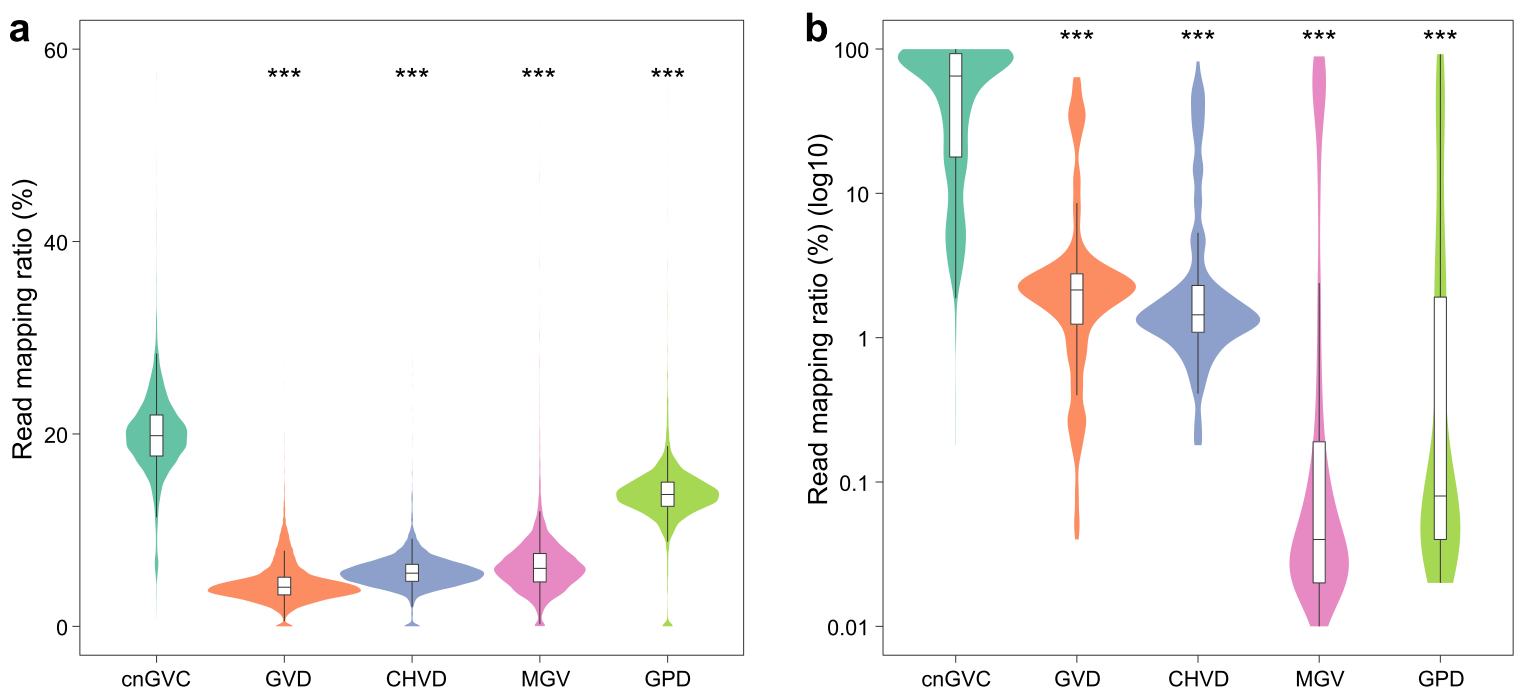

**Supplementary Fig. 6| Comparison of read mapping ratio between cnGVC and other gut viral databases. a-b,** The read mapping ratio of bulk metagenome samples **(a)** and VLP-based metagenome samples **(b)**. For **b**, the values were shown using a logarithmic coordinate. Wilcoxon rank-sum test: *, *p*<0.05; **, *p*<0.01; ***, *p*<0.001.

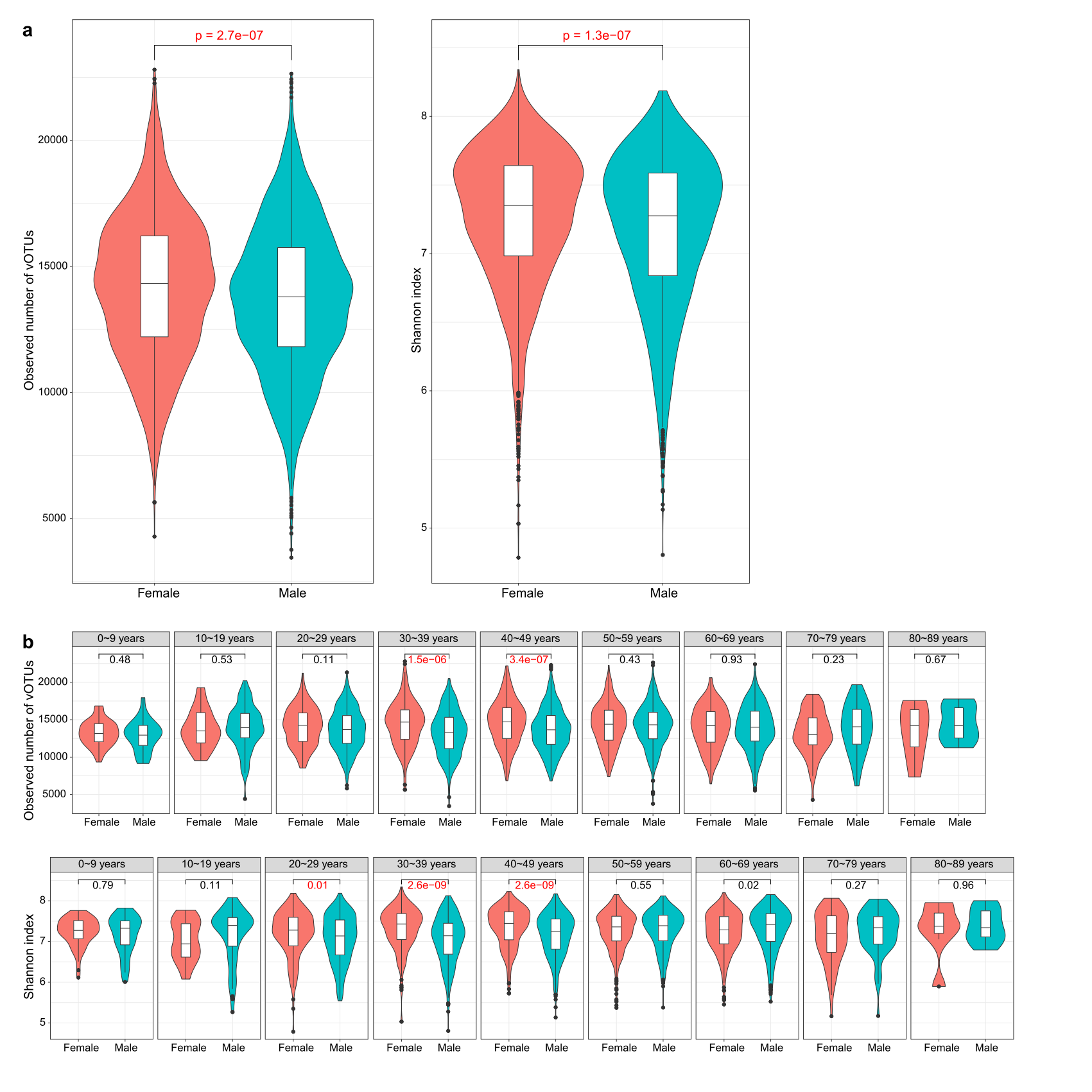

**Supplementary Fig. 7| Sex-related variations in the richness and diversity of the gut virome. a,** Comparison of gut virome richness and diversity between females and males. **b,** Comparison of gut virome richness and diversity between females and males grouped by age stages. For **b**, the statistical test was performed using Wilcoxon rank-sum test and the q-values were shown. Boxes represent the interquartile range between the first and third quartiles and the median (internal line). Whiskers denote the lowest and highest values within 1.5 times the range of the first and third quartiles, respectively; dots represent outlier samples beyond the whiskers.

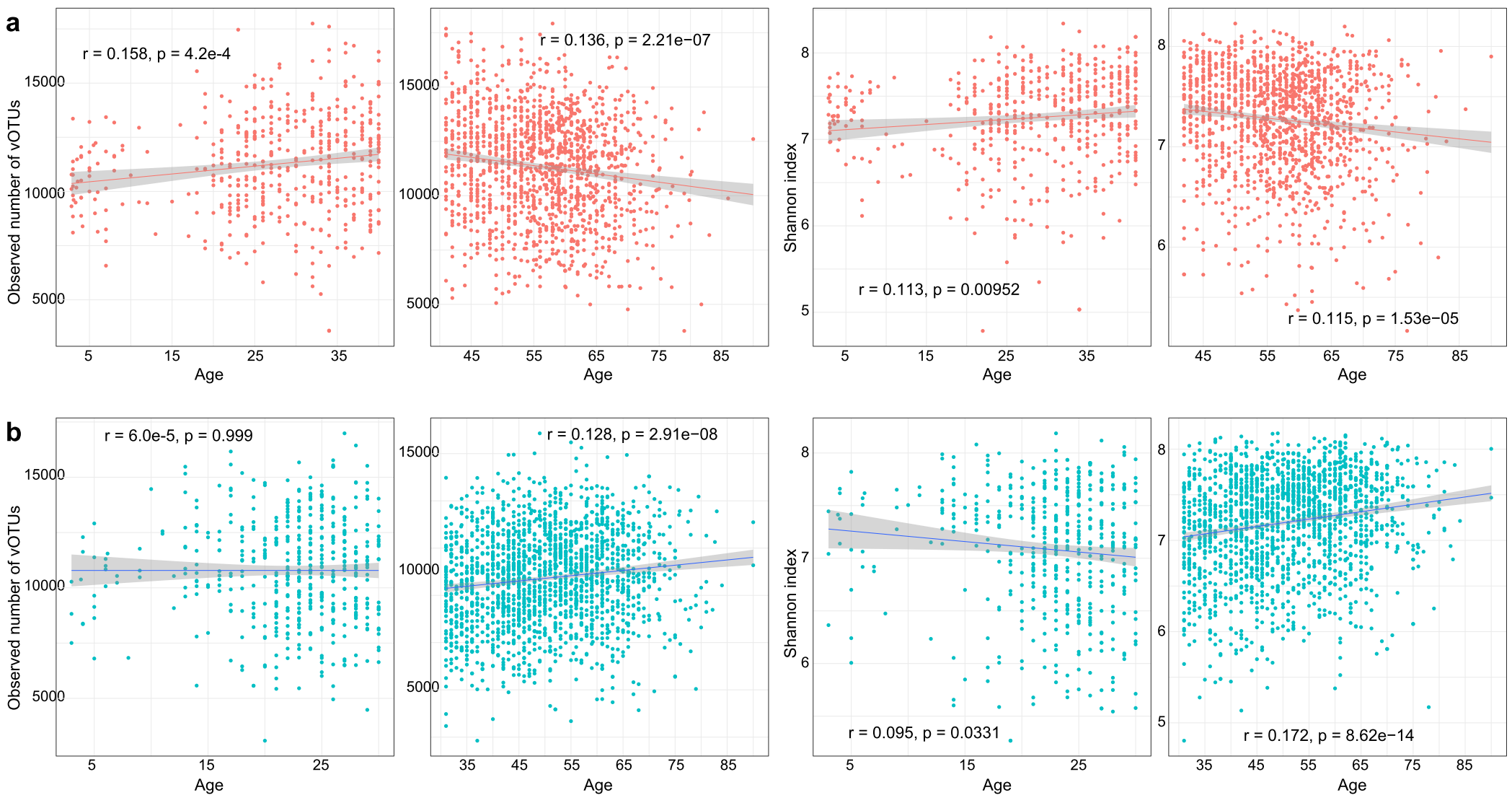

**Supplementary Fig. 8| Age-related variations in the richness and diversity of the gut virome. a-b,** Scatter plots showing age-related variations of virome richness (left two panels) and diversity (right two panels) for females **(a)** and males **(b)**. For females, values were split at the age of 40. For males, values were split at the age of 30. The Pearson correlation coefficients *r* and the p-values are shown.

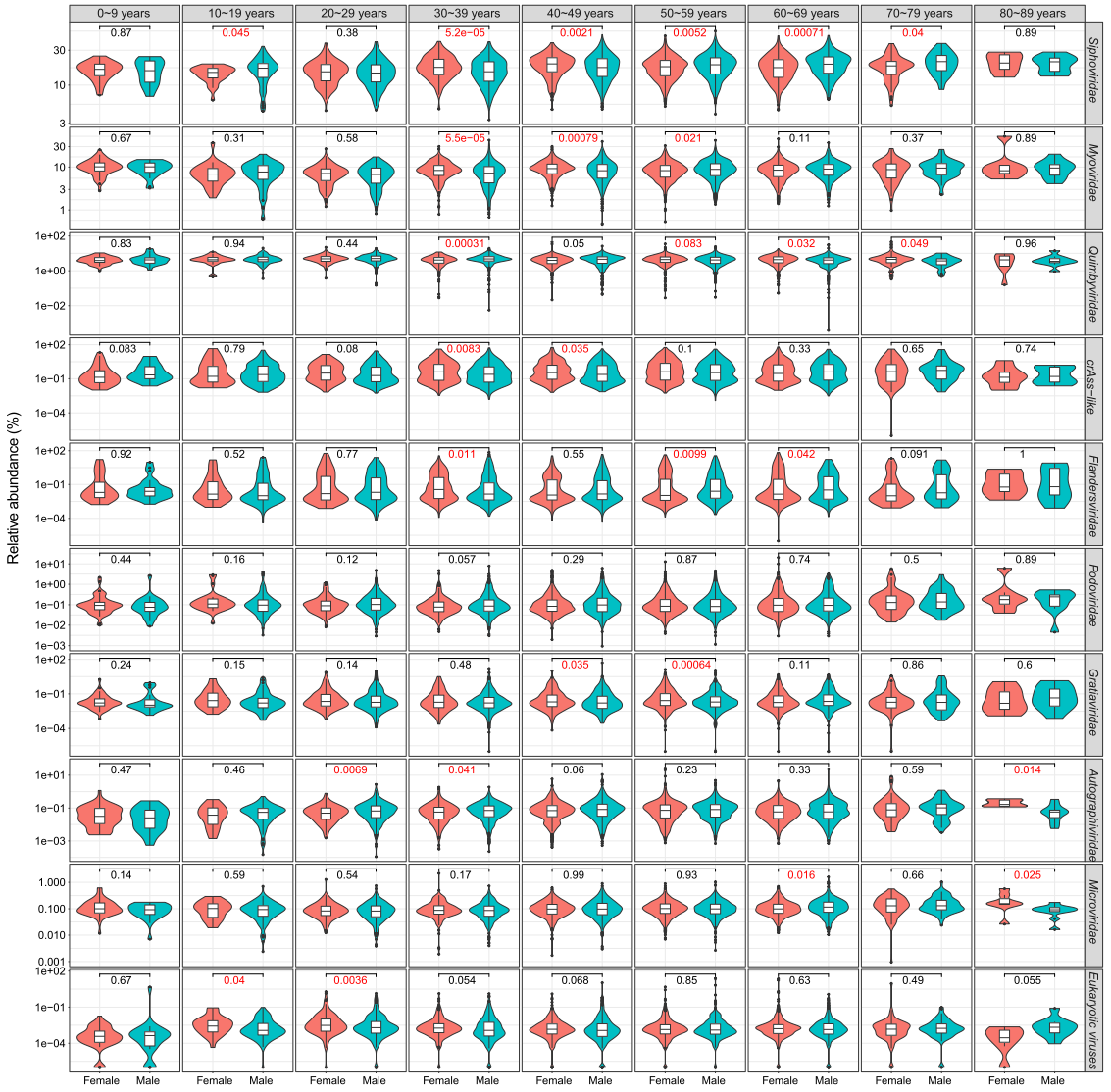

**Supplementary Fig. 9| Sex and age-related variations of the gut virome at the family level.** Statistical test was performed using Wilcoxon rank-sum test and the q-values were shown. Boxes represent the interquartile range between the first and third quartiles and the median (internal line). Whiskers denote the lowest and highest values within 1.5 times the range of the first and third quartiles, respectively; dots represent outlier samples beyond the whiskers.

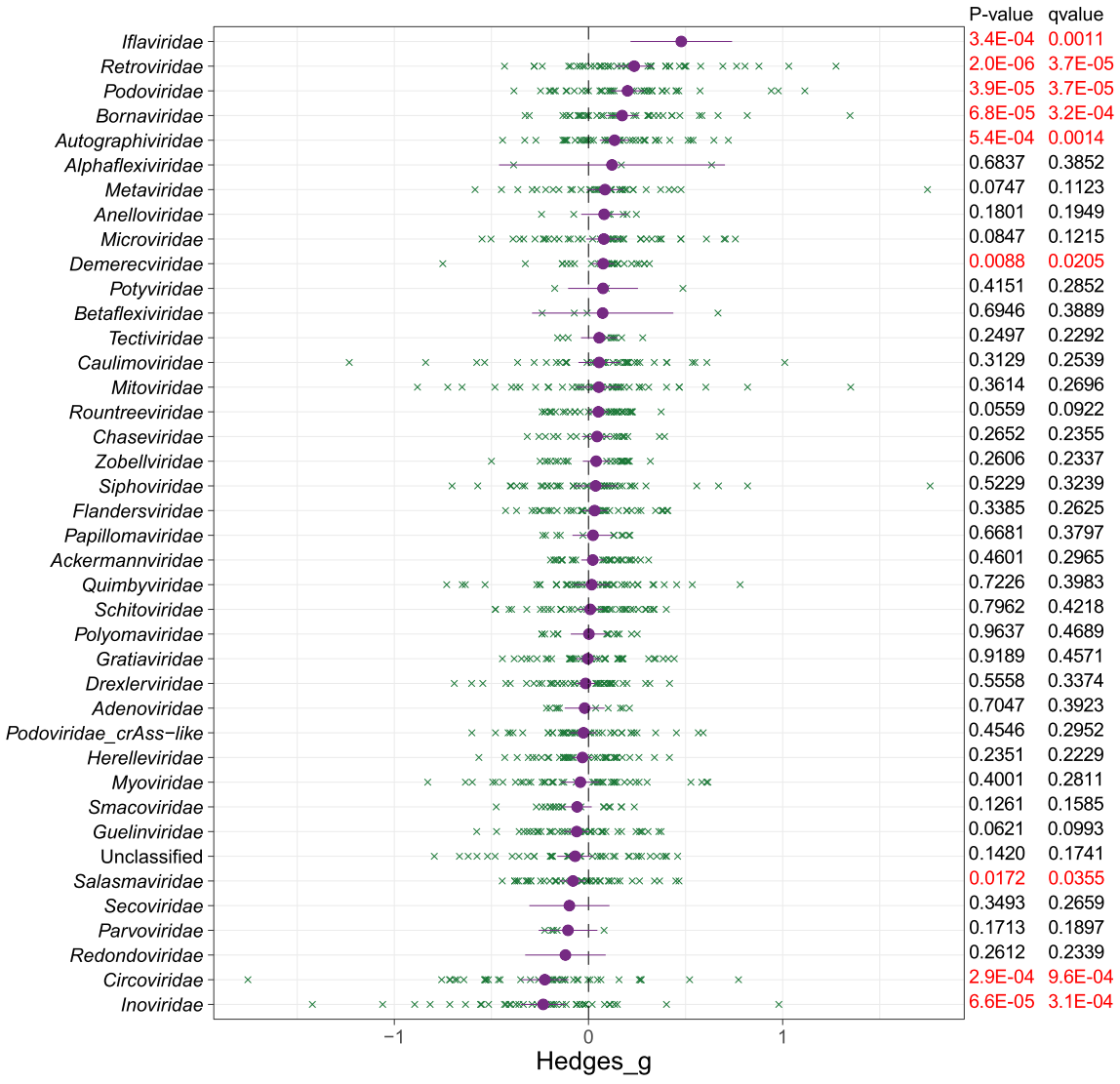

**Supplementary Fig. 10| Meta-analysis of the abundances of viral families across common diseases.** Forest plot showing the Hedges' g standardized mean differences and the random effects model based on the arcsine-square root-transformed abundances of viral families. Case-control comparisons are labeled by green crosses. Solid black lines indicate the 95% confidence intervals. The meta-analysis p-values and q-values are shown in the right panel.

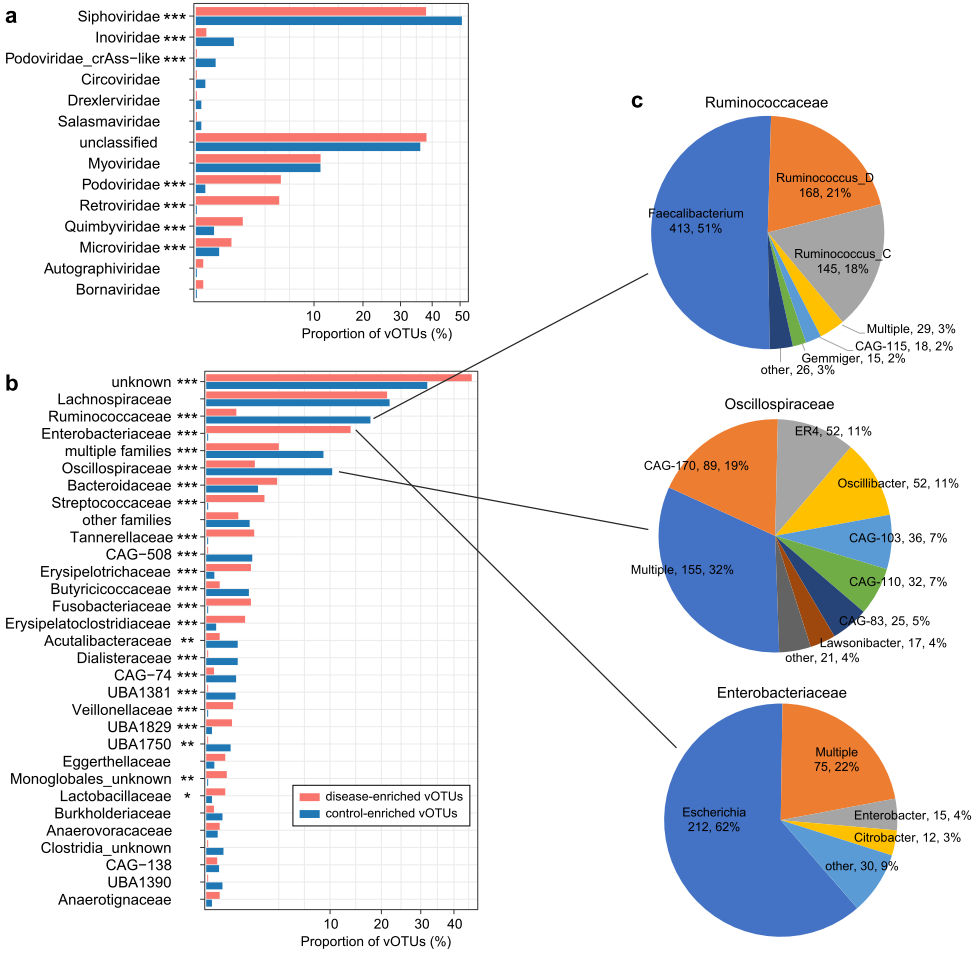

**Supplementary Fig. 11| Taxonomy and host assignment of the viral signatures. a-b**, Comparison of taxonomic annotation **(a)** and host assignment **(b)** between the disease-enriched and control-enriched vOTUs. The vOTUs are grouped at the family level, and the prokaryotic host taxa are also shown at the family level. **c,** Pie plot showing the genus-level host assignment of vOTUs that are predicted to infect Ruminococcaceae, Oscillospiraceae, and Enterobacteriaceae. Fisher’s exact test: *, *q*<0.05; **, *q*<0.01; ***, *q*<0.001.

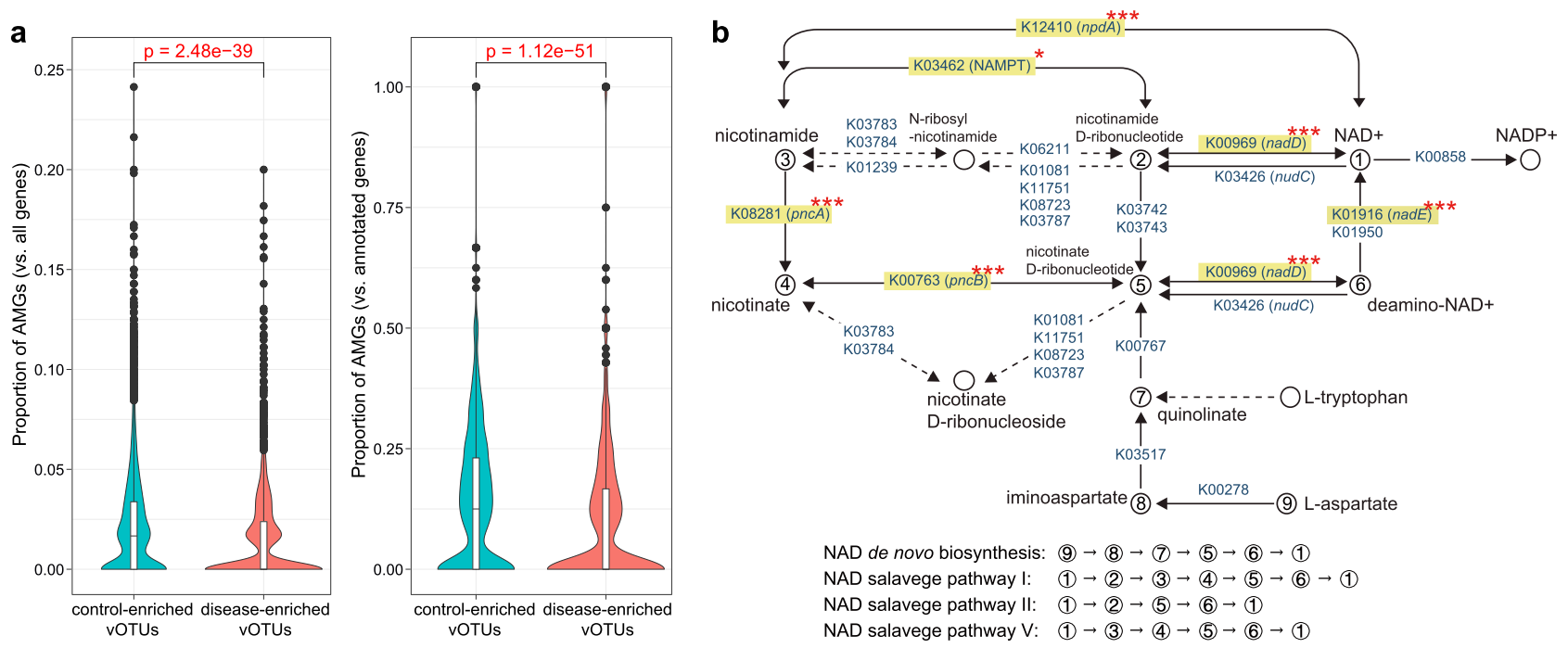

**Supplementary Fig. 12| Functions of the disease-associated viral signatures. a,** Boxplots showing the proportions of auxiliary metabolic genes (AMGs) in disease-enriched and control-enriched vOTUs. Boxes represent the interquartile range between the first and third quartiles and the median (internal line). Whiskers denote the lowest and highest values within 1.5 times the range of the first and third quartiles, respectively; dots represent outlier samples beyond the whiskers. **b,** Enzymes and pathways involving nicotinamide adenine dinucleotide (NAD+) *de novo* biosynthesis and salvage. This diagram is exacted from the KEGG pathway database with manual modifications. Six key enzymes with a higher frequency in the control-enriched vOTUs are labeled by yellow boxes. Fisher’s exact test: *, *q*<0.05; **, *q*<0.01; ***, *q*<0.001.

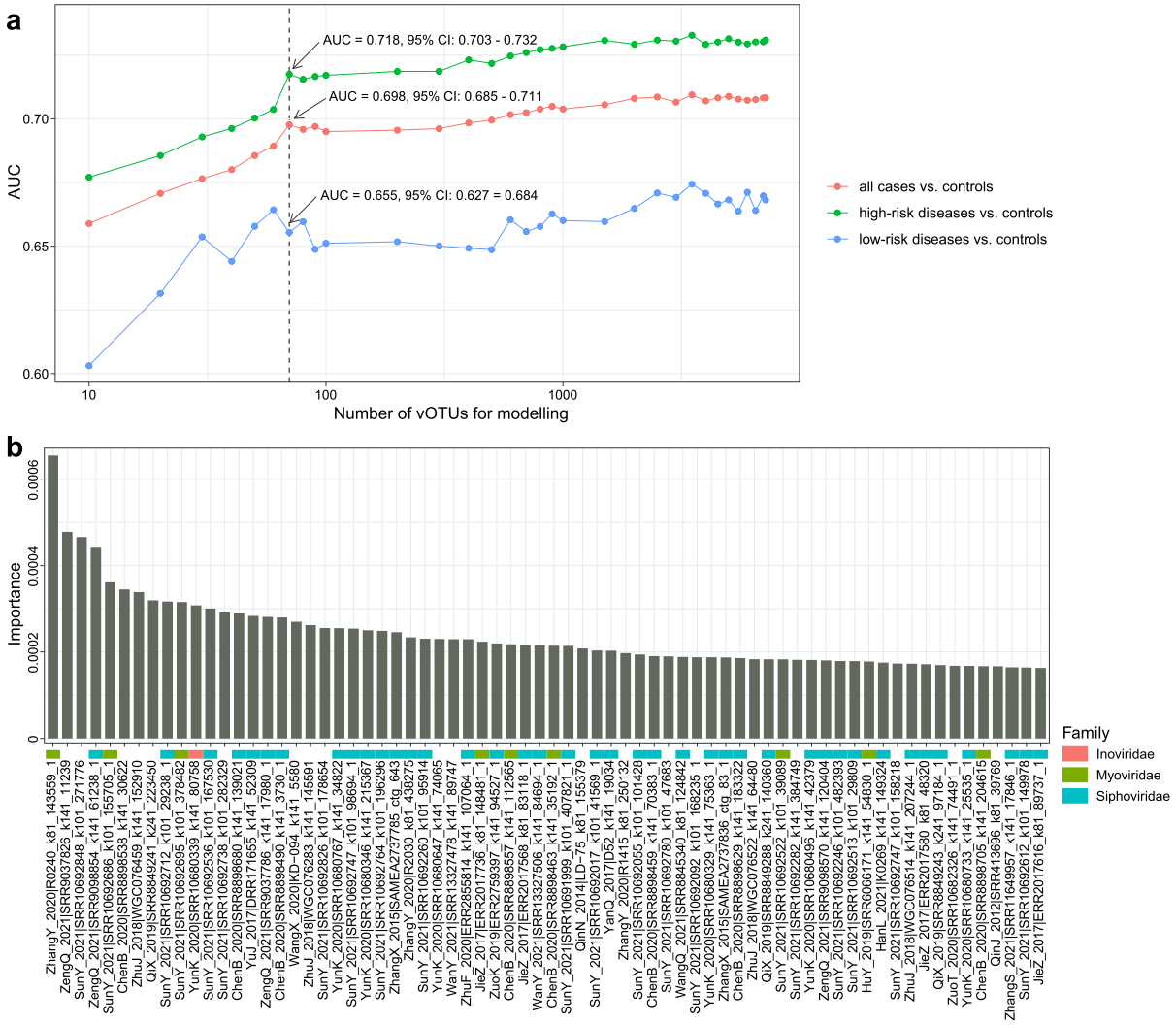

**Supplementary Fig. 13| Random Forest models for discriminating patients and controls using the universal viral signatures. a,** The performance is explored for different numbers of species, ordered in importance. Classification performance of a random forest model assessed by area under the receiver-operating characteristic curve (AUC). This analysis shows that the 70 most important vOTUs generated the discriminatory power of AUC 0.698 (95% CI, 0.685-0.711) for classifying all samples and 0.718 (0.703-0.732) for high-risk diseases versus controls. **b,** Detailed information of the 70 most important vOTUs.

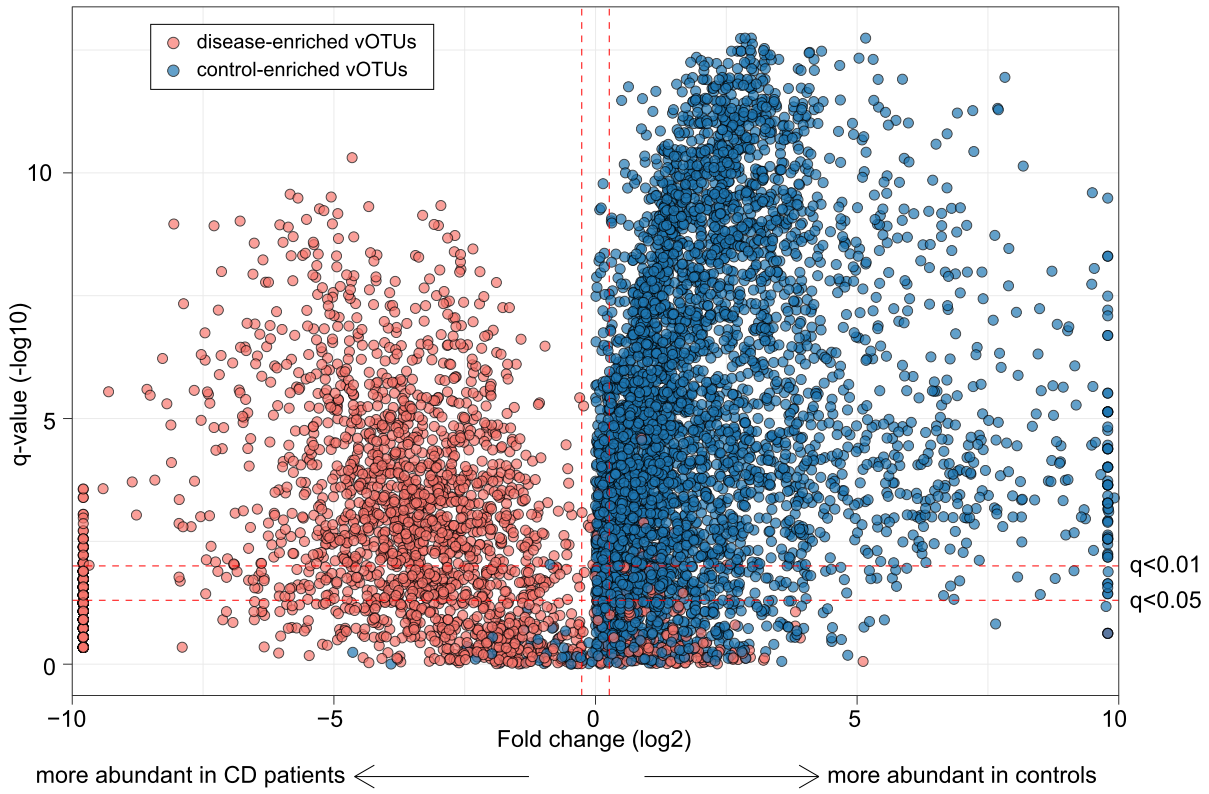

**Supplementary Fig. 14| Alterations of universal viral signatures in Crohn's disease.**

Volcano plots showing the fold change vs. q-values for 7,161 disease-associated vOTUs in the Crohn’s disease (CD) cohort. The X-axis shows the ratio (log2 transformed) of vOTU abundance in disease cases (fold<0) compared with that in healthy controls (fold>0). The Y-axis shows the q-value (-log10 transformed) of a vOTU. vOTUs are colored by their enrichment directions in CKD patients vs. healthy controls. Horizontal dotted lines: *q*<0.05 and *q*<0.01; vertical dotted lines: fold<-1.2 and fold>1.2.
